# Area-level context and functional brain organization across the lifespan

**DOI:** 10.64898/2026.07.21.739899

**Authors:** Jocelyn A. Ricard, Chase Antonacci, Gabriel Reyes, Eugenia Giampetruzzi, Jivesh Ramduny, Gracie Grimsrud, Monica Ellwood-Lowe, Russell A. Poldrack

**Author notes:** **Corresponding Author:** Jocelyn A. Ricard, 450 Jane Stanford Way, Building 420, Stanford University, Stanford, CA 94305.

## Abstract

The neighborhoods children grow up in are associated with cognitive and mental health outcomes across the lifespan. Yet how neighborhood-level factors are quantified, and how they relate to large-scale functional brain connectivity, remains an emerging area of investigation. This scoping review synthesizes empirical evidence on associations between area-level factors and resting-state functional brain connectivity and network organization. We included studies assessing resting-state fMRI functional connectivity or rs-fMRI-derived intrinsic activity in relation to validated area-level deprivation indices guided by Trinidad et al.’s (2022) inventory, the Opportunity Atlas, or geocoded crime exposure from the Uniform Crime Reporting program. Of the 28 studies meeting inclusion criteria, the evidence base was unevenly distributed across the lifespan; 25 of 28 (89.3%) examined perinatal, childhood, or adolescent samples, while only one examined adults exclusively. The literature was also methodologically non-independent, over half drew on the Adolescent Brain and Cognitive Development (ABCD) Study, and the Area Deprivation Index was the most commonly used measure (k = 16, 57.1%). Therefore, apparent convergence partly reflects shared data and instruments rather than independent replication. Where directions could be compared, higher area-level disadvantage was associated predominantly with lower within-network connectivity in association and control systems and higher connectivity in somatomotor systems, with several findings attenuating once area-level exposure was isolated from household socioeconomic status. Geographic resolution varied but was rarely justified. Progress will require greater theoretical intentionality in selecting area-level measures, transparent justification of geographic resolution, treatment of dataset and measure non-independence, and investment in understudied developmental periods and non-US contexts.

## Introduction

Neighborhoods are a foundational context for human development^1–4^. In research, neighborhood context is typically captured through area-level measures, i.e., census-derived indices that aggregate conditions at a macro-level such as poverty, crime, and resource access across defined geographical units. Where people live shapes their access to quality schools, safe green spaces, healthy food, employment, and healthcare, resources that are unequally distributed across communities in ways that track closely with race, class, and historical policy^5–7^. In the United States, concentrated neighborhood disadvantage disproportionately affects Black, Indigenous, and other racially marginalized communities, reflecting decades of contextual policies including residential segregation, redlining, and disinvestment^8,9^. Area-level factors are those that reflect properties of a particular environment or spatial location at a particular moment in time that are aggregated across a community of people (e.g., neighborhood or county-level measures), and such factors as poverty, violence, or segregation have been shown to have important implications for development independent of individual and family characteristics^3,10–12^.

How these area-level neighborhood factors may become biologically embedded in the brain over time and the neural mechanisms through which this embedding may occur, however, are not well characterized. Understanding how area-level disadvantage shapes brain development is therefore a pressing scientific and public health question. The brain represents a central biological mechanism through which environmental conditions may be transduced into long-term developmental outcomes ^11,13^. Stress-sensitization frameworks suggest that chronic exposure to resource-poor, high-threat environments shapes neuroendocrine, autonomic, and neural systems in ways that reflect adaptation to adversity, with potential costs to cognitive and affective regulation ^14,15^. Functional neuroimaging may therefore provide a powerful window into these processes, enabling researchers to characterize how area-based exposures alter brain organization across development. This aligns with a recent call for a paradigm shift in cognitive neuroscience to incorporate upstream macro-social contextual factors when investigating the neural correlates of macrosocial inequity ^9^. In particular, given that resting-state functional connectivity and network topography represent the temporal coordination of spontaneous neural activity across large-scale functional networks ^16,17^, functional connectivity may therefore capture more proximally responsive or dynamic aspects of area-level impacts than measures of brain structure alone ^18^.

Yet several open questions limit what can be concluded from this emerging work. Prior work has operationalized socioeconomic deprivation in neuroscience using an array of measures across individual-, household-, and neighborhood-level factors, while often used interchangeably under the umbrella of “socioeconomic status” ^13,19^. This heterogeneity extends to how resting-state function itself is operationalized, from edge-level connectivity to whole-brain network organization. It is also important to consider their distinct dimensions across the lifespan, as the neural correlates of area-level disadvantage are unlikely to be uniformly distributed across development. Early childhood represents a sensitive period for corticolimbic circuitry supporting threat and stress regulation, while adolescence involves protracted maturation of association cortices along a sensorimotor-to-association axis, with higher order networks at the end of this hierarchy showing heightened sensitivity to environmental input^20,21^. Lastly, older age may reflect the accumulated burden of chronic area-level exposures^22,23^. Attending to developmental timing is therefore important for identifying when area-level context exerts its greatest influence on functional brain organization.

Together, these open questions, spanning how area-level context and brain function are each operationalized, how effects may vary across development, and whether they generalize across contexts, point to a need to take stock of what has been studied and where gaps and misalignment remain. Therefore, this scoping review aims to synthesize the current literature on the relationship between area-level factors and functional brain connectivity and network topology across the lifespan.

### Area-Level Measures for Understanding the Brain

A critical challenge in this literature is the conceptual and methodological heterogeneity of how area-level factors are operationalized. Prior neuroscience research has frequently conflated individual-level socioeconomic status (e.g., household income, parental education) with area-level factors, despite evidence that these constructs capture distinct dimensions of inequality with potentially distinct implications for brain development ^12,19^. Area-level measures capture the collective conditions of a place, independent of any one person’s resources, and are typically aggregated at the census-block, tract, ZIP code, or county level. A recent scoping review of area-based deprivation measures used in U.S. health research identified over a dozen validated composite indices, including the Area Deprivation Index (ADI; ^24,25^), Social Vulnerability Index (SVI; ^26^), and Child Opportunity Index (COI; ^27,28^), among others ^29^. Trinidad et al. (2022) surveyed area-level socioeconomic indices used in United States health research; here we adopt their inventory as a starting point, while noting that this scope carries implications for the geographic generalizability of findings, which we return to in the Discussion. These indices differ meaningfully in their construction, domains, and target populations, see Table 1 for full list of indices used. No prior review has systematically synthesized this literature across the lifespan.

**Table 1.**
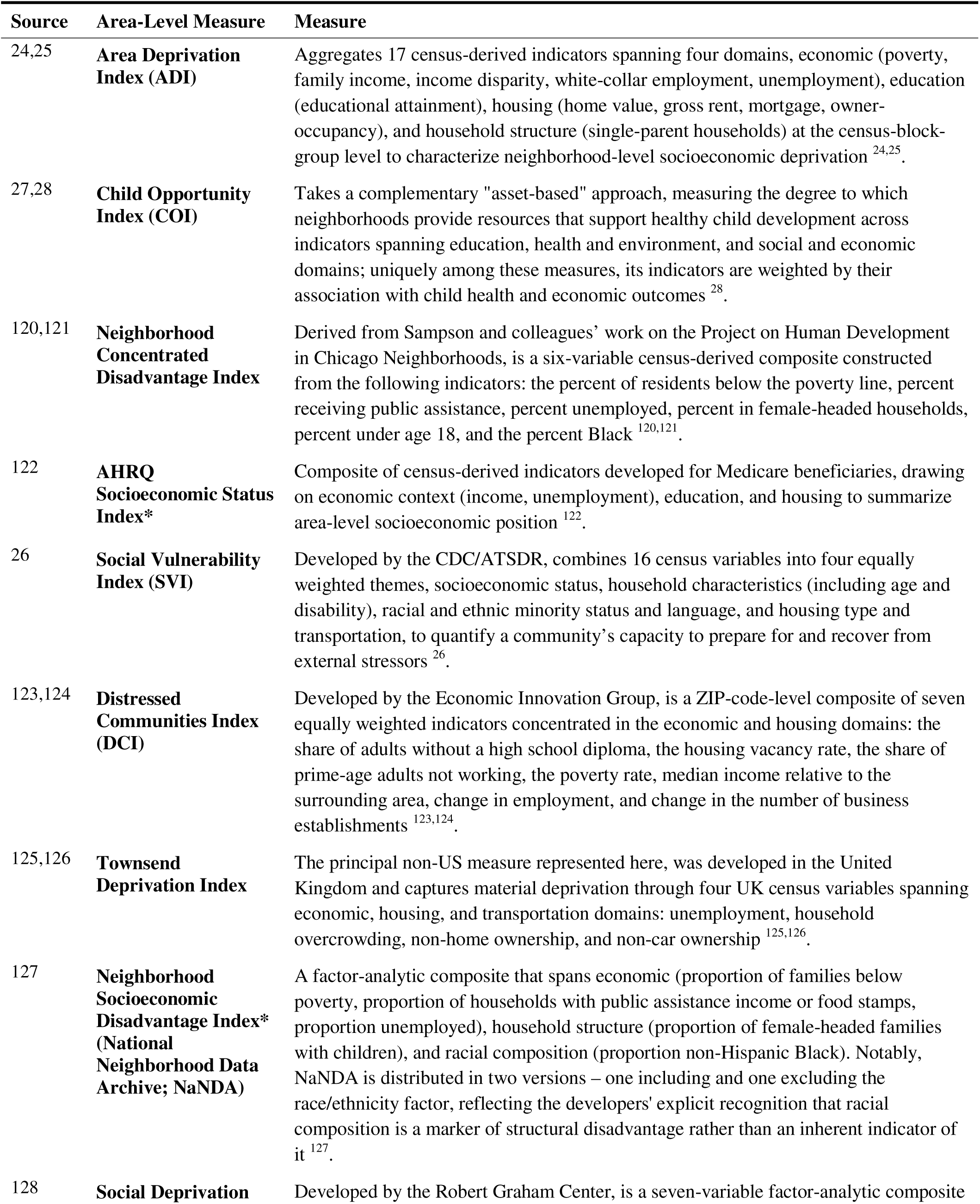
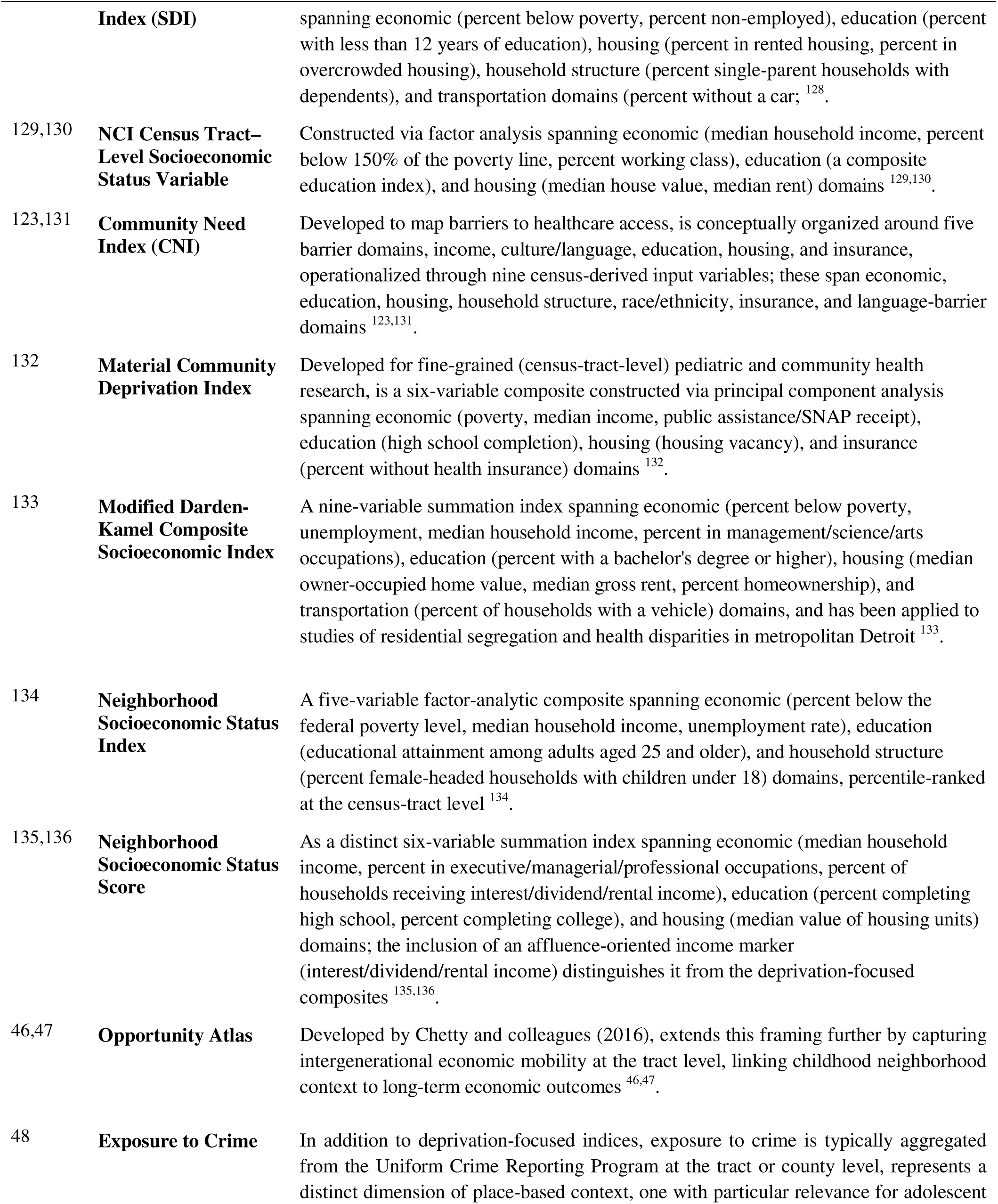
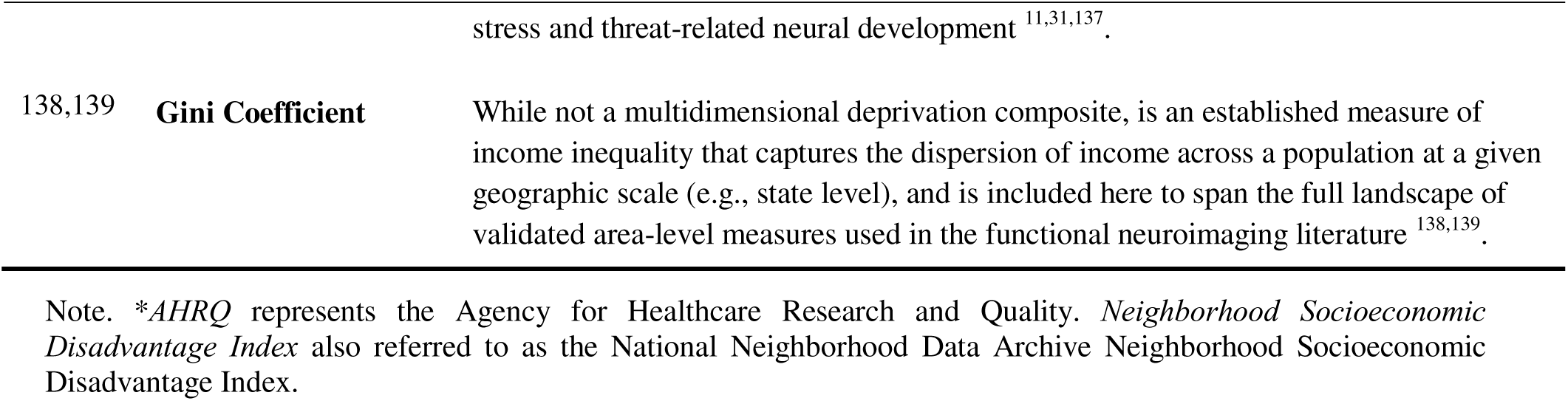
Eligible area-level deprivation and opportunity indices. Glossary of the validated composite area-level measures eligible for this review. For each measure the table lists the source reference, the index name, and a description of its construction, constituent domains, and native geographic unit.

Despite the availability of these validated instruments, their uptake in neuroimaging research has been uneven. Many studies rely on multidimensional composite indices like the Area Deprivation Index, but the choice of index is rarely theoretically motivated. The ADI is freely available through a well-established public interface ^25^ and, critically, is the area-level measure pre-linked to residential addresses in the ABCD Study, the dataset underlying half of the studies reviewed here. The dominance of the ADI may therefore reflect data infrastructure and convenience as much as deliberate construct selection. Recent efforts to expand the environmental measures linked to large neuroimaging cohorts have similarly noted the field’s disproportionate reliance on a narrow set of deficit-oriented socioeconomic composites, and have called for incorporating complementary asset-based and multidimensional place-based indicators ^30^. This is important because the ADI captures a specific dimension of area-level context, material socioeconomic deprivation, and does not index the presence of developmental resources, exposure to threat and violence, or other dimensions of vulnerability. Complementary measures are designed to capture what the ADI omits, e.g., geocoded crime and violence exposure indexes threat as a dimension theoretically distinct from deprivation ^31^, and multidimensional indices such as the Social Vulnerability Index span housing, disability, and language domains orthogonal to economic hardship. However, these measures remain less used in the neuroimaging literature corpus.

A growing body of neuroimaging research has investigated associations between neighborhood disadvantage and brain structure. A recent systematic review identified 37 studies examining neighborhood conditions and structural brain outcomes in children and adolescents, finding that adverse neighborhood socioeconomic conditions were linked to reduced brain volume and white matter integrity, and smaller cortical surface areas ^32^. Notably, observed racial disparities in brain structure were partially explained by residence in low-resourced neighborhoods ^33,34^. Importantly, in the United States, these neighborhood conditions reflect a history of racialized disinvestment, so such disparities likely index differential exposure to structural conditions rather than inherent biological differences between racial groups ^35^. In particular, mediation analyses across multiple studies found relationships between race and regional brain volume and cortical thickness were significantly mediated by neighborhood disadvantage ^33,34^, underscoring the role of neighborhood context in shaping neurodevelopment. Yet the structural neuroimaging literature, while informative, offers only a partial view of how neighborhoods shape the brain. Structural MRI captures morphological properties of brain tissue, e.g., white matter, cortical thickness, and/or volume, that develop and change on slow timescales, and are thought to reflect the cumulative, longer-term consequences of exposures. In contrast, functional connectivity measured during resting-state fMRI (rs-fMRI) indexes the dynamic, large-scale coordination among brain regions that underlies cognition, emotion, and behavior. Furthermore, resting-state functional connectivity provides a systems-level perspective, enabling the characterization of whole-brain network features, e.g., modularity or integration, that are not captured by structural metrics alone.

### Resting-State Functional Connectivity as an Outcome

Resting-state fMRI (rs-fMRI) measures spontaneous fluctuations in blood oxygen level dependent (BOLD) signal in the absence of an explicit task, and functional connectivity refers to the statistical dependence of these fluctuations between brain regions over time ^36,37^. The organization of the brain into large-scale intrinsic networks, including the default network (DMN), salience network (SN), and frontoparietal/executive control network (FPN), is highly reproducible across individuals and is sensitive to both developmental change and environmental variation ^38,39^. These networks support the kinds of cognitive and affective functions, such as self-referential processing, threat detection, and cognitive control, that are most likely to be shaped by chronic neighborhood adversity. Emerging evidence suggests that area-level factors are associated with altered functional connectivity in children and adolescents. For example, neighborhood poverty during childhood has been prospectively linked to reduced functional network segregation and integration balance during adolescence, with effects concentrated in early but not later adolescence ^40^. However, this literature remains fragmented given studies vary widely in the area-level measures employed, the functional connectivity outcomes assessed, the age ranges examined, and the degree to which individual- and family-level socioeconomic factors are accounted for.

The goal of this scoping review is to map and synthesize empirical evidence on associations between area-level factors and resting-state functional brain connectivity and network organization across the lifespan. Guided by the JBI methodology for scoping reviews ^41^ and the PRISMA-ScR reporting standards ^42^, we systematically searched four databases for studies that used rs-fMRI to examine functional connectivity in relation to validated area-level deprivation indices, crime exposure, or the Opportunity Atlas. Unlike prior reviews focused on structural brain outcomes or exclusively on periods of development ^32,43^, this review spans the full lifespan and is specifically focused on functional connectivity as a distinct outcome.

This review makes several contributions to the literature. First, it provides a comprehensive synthesis of the functional neuroimaging literature on area-level factors, distinguishing these from individual- and household-level SES measures. Second, by mapping the range of area-level measures and functional connectivity outcomes used across studies, it identifies methodological patterns and gaps that can inform the design of future research. Third, by adopting a lifespan framing, it enables consideration of developmental specificity in how neighborhood context shapes functional brain organization, a question with important implications for sensitive periods of intervention. Together, this review responds to a growing call for neuroscience to engage with the upstream contextual determinants of brain health ^9,44,45^, and provides a review base to guide the field forward.

## Methods

This scoping review was conducted in accordance with the Joanna Briggs Institute (JBI) methodology for scoping reviews ^41^ and the Preferred Reporting Items for Systematic Reviews and Meta-Analysis Extension for Scoping Reviews (PRISMA-ScR) Checklist and Explanation document ^42^. The review protocol was preregistered with the Open Science Framework on September 29th, 2025 (https://doi.org/10.17605/OSF.IO/S7VE6). Any significant amendments to the pre-registered protocol are detailed and published alongside the completed scoping review.

### Review Objectives

This scoping review was guided by the following primary research question: What is known about the relationship between area-level neighborhood factors and large-scale functional brain connectivity and network organization across the lifespan?

### Search Strategy

A systematic search was conducted across four electronic databases: PubMed, Embase, Web of Science, and SCOPUS. Searches were conducted with no restrictions on the date of publication, capturing the full historical literature on the topic. Search terms combined controlled vocabulary and free-text terms spanning three conceptual domains: (1) brain and functional connectivity (e.g., functional connectivity, resting state, functional networks), (2) area-level contextual and neighborhood characteristics (e.g., neighborhood deprivation, neighborhood disadvantage, community violence, exposure to crime), and (3) validated composite area-level deprivation indices (e.g., Area Deprivation Index, Child Opportunity Index, Social Vulnerability Index, Opportunity Atlas). Searches were conducted on October 6-7, 2025 and restricted to English-language publications, and editorial and review article publication types were excluded at the search stage. In addition to the four electronic databases, we conducted supplementary hand-searching in Google Scholar to capture relevant records not indexed by the primary databases; this yielded two additional records, which were screened against the same eligibility criteria. The full search string as implemented in each database is provided in the Supplementary Materials. Consistent with the scope expansions described in Deviations from Preregistered Protocol, we interpreted resting-state functional connectivity to encompass rs-fMRI-derived measures of intrinsic functional brain organization, including edge-level, within- and between-network, and graph-theoretic connectivity.

## Eligibility Criteria

Eligibility criteria were defined in accordance with JBI recommendations for scoping reviews.

### Inclusion criteria

Studies were included if they met all of the following criteria:

- Participants: Human participants of any age (children through older adults), reflecting the lifespan scope of this review.
- Employed functional magnetic resonance imaging (fMRI) to assess resting-state functional connectivity, including edge-level connectivity between specific regions of interest, within- and between-network connectivity among large-scale brain networks (e.g., default mode, salience, executive control networks), and/or graph-theoretic measures of network organization (e.g., modularity).
- Assessed area-level factors using validated composite indices of deprivation or complementary place-based measures. Eligible indices were guided by Trinidad et al.’s (2022) scoping review of commonly used area-level socioeconomic deprivation measures in U.S. health research, and additionally included the Opportunity Atlas ^46,47^ and county-level exposure to crime aggregated from the Uniform Crime Reporting Program ^48^.
- Studies conducted in any country or setting.
- Peer-reviewed published journal articles available in English.

### Exclusion criteria

Studies were excluded if they met any of the following criteria:

- Non-empirical research papers, including reviews, book chapters, commentaries, perspectives, or opinion pieces.
- Studies using neuroimaging modalities other than fMRI (e.g., structural MRI, diffusion MRI, Positron Emission Tomography [PET], electroencephalography [EEG], functional near-infrared spectroscopy [fNIRS]).
- Studies that exclusively used task-based fMRI or did not report a measure of functional connectivity.
- Non-peer reviewed or non-published materials (e.g., theses/dissertations, protocol papers, preprints, retracted papers, conference abstracts, popular-science books).

### Area-Level Measures

Initial selection of eligible area-level deprivation indices was guided by Trinidad et al.’s (2022) scoping review of commonly used area-level socioeconomic deprivation measures in U.S. health research. Eligible indices capture neighborhood resources and conditions aggregated at the census-tract or county level and are listed in Table 1. While the Gini coefficient was not part of the original preregistration, Trinidad et al. 2022, identify it as an established measure of income inequality, distinct from, but adjacent to multidimensional deprivation composites. We include it here to capture the broad landscape of validated area-level neighborhood measures used in functional neuroimaging research. Because the Trinidad et al. (2022) inventory was developed within United States health research, our eligible exposures list primarily reflects US-originated indices, with the Townsend Deprivation Index as a notable non-US exception. Parallel instruments developed in other national contexts (e.g., the English Indices of Multiple Deprivation) were not separately targeted by the search strategy in the scope of this review. Several included studies operationalized area-level neighborhood context through single census-derived indicators (e.g., tract-level neighborhood poverty rate; tract-level family poverty rate) rather than validated multidimensional composites. These were retained where the exposure was clearly aggregated at an area level (block group, tract, ZIP, county, or state) rather than at the individual or household level, on the grounds that the area-level character of the construct, not the multidimensional construction of the index, is the central inclusion criterion of this review.

### Study Selection

Upon completion of the database searches, all identified citations were collated and uploaded into Covidence (https://www.covidence.org/) for de-duplication and screening. Titles and abstracts were independently screened by two reviewers against the eligibility criteria outlined above. The full text of all potentially eligible citations was subsequently assessed in detail by two independent reviewers. Reasons for exclusion were documented at each stage of the screening process. Disagreements arising between reviewers at any stage were resolved through group discussion or by consulting an additional reviewer. This process resulted in 28 articles meeting inclusion criteria, and the full PRISMA-ScR flow diagram is reported in **Figure 1**.

**Figure 1:**
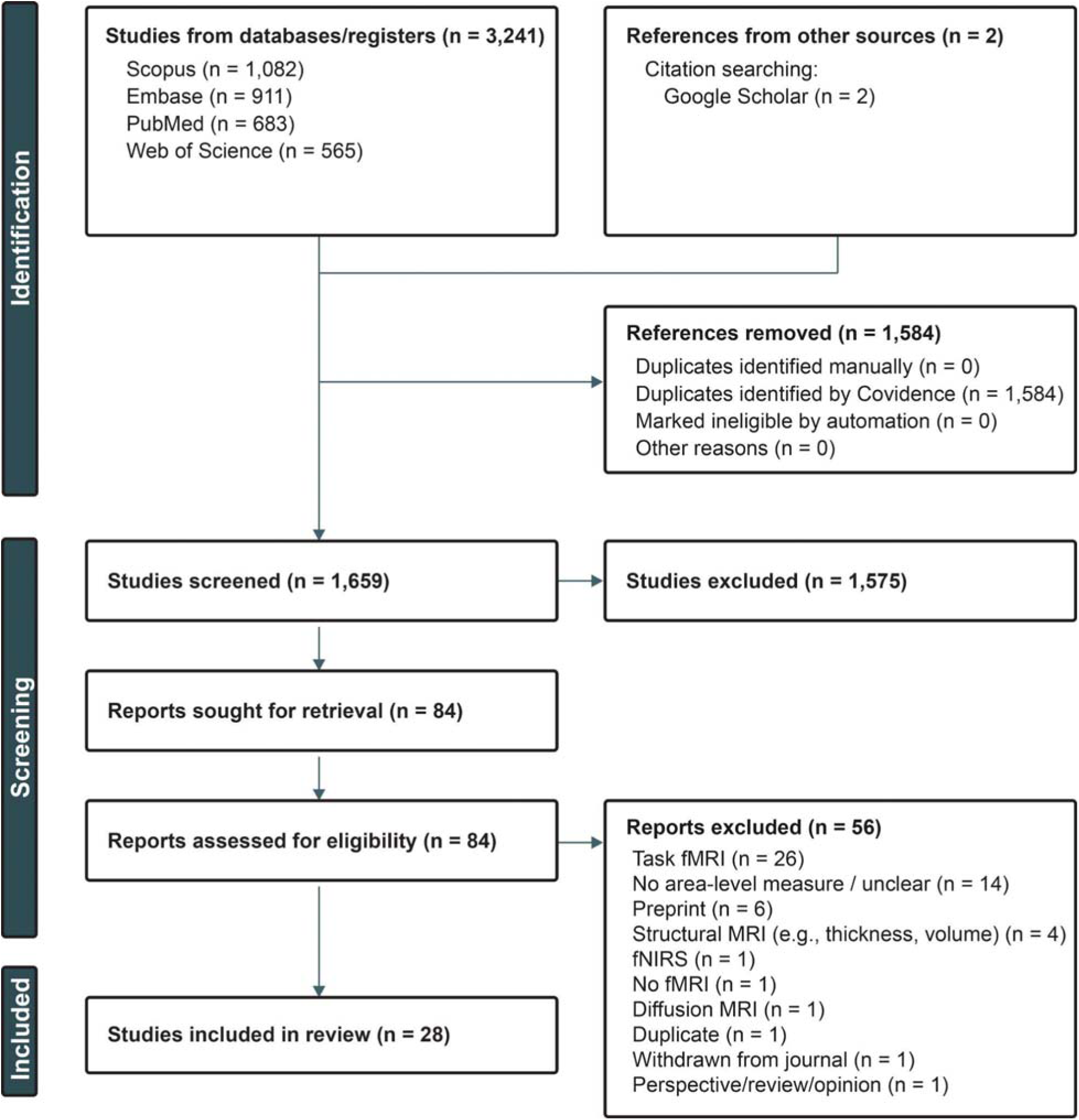
PRISMA Flow Diagram. The figure summarizes record identification, screening, eligibility assessment, and final inclusion, with exclusions documented at each stage based on predefined criteria.

## Data Extraction

Data were extracted from all included articles using a standardized extraction form developed by the review team and piloted prior to full extraction. Extractions were reviewed by a second independent reviewer for consistency and accuracy. The following variables were extracted from each included study:

- Study and publication year
- Total sample size
- Area-level neighborhood measure(s) used
- Covariates included in analyses
- Sociodemographic characteristics of the sample (e.g., age, race/ethnicity)
- Neuroimaging measure of functional connectivity (e.g., seed-based connectivity, ROI-to-ROI, independent component analysis/resting-state networks, whole-brain, graph-theoretic metrics)
- Brain parcellation scheme, where applicable
- Study design characteristics (e.g., cross-sectional, longitudinal, predictive modeling)
- Direction and summary of primary findings (positive, negative, or null associations)

### Data Synthesis

Consistent with JBI scoping review methodology and the heterogeneous nature of the included literature, findings were synthesized descriptively and narratively rather than via meta-analytic pooling. Extracted data were organized and tabulated to map the scope of the existing literature, characterize the area-level measures and neuroimaging approaches employed, and identify convergent and divergent patterns in findings across developmental periods (childhood, adolescence, adulthood, and older adulthood). Particular attention was paid to the diversity of area-level measures used, the functional connectivity outcomes assessed, and gaps in the literature warranting future investigation.

### Deviations from Preregistered Protocol

The registered protocol specified validated composite indices of deprivation (guided by Trinidad et al., 2022), the Opportunity Atlas, and UCR-based crime exposure as eligible area-level exposures. During the screening, four additional categories were retained that extended beyond the initial scope.

1. Several studies operationalized area-level context through a single census-derived variable (e.g., tract-level family poverty rate; ^40^) rather than a multidimensional composite index. These were retained because the inclusion criterion (e.g., assessed area-based factors) prioritized the area-level character of the exposure over the specific construction of the index. We acknowledge this expands the independent variable definition specified in the protocol.
2. Macro-scale socioeconomic indices, i.e., the state-level Gini coefficient ^49^, were retained because Trinidad et al., 2022 identify it as an established economic inequality measure adjacent to multidimensional deprivation composites, though it was not listed in the preregistered table of indices. We note that state-level operationalization captures a substantively different construct than block-group or tract-level deprivation composites.
3. Parcel-wise measures of regional intrinsic activity (e.g., BOLD fluctuation amplitude, ALFF/fALFF, ReHo) were retained because they derive from the same rs-fMRI signal and capture organizationally meaningful properties, as done in Sydnor et al., 2023, even though they are not connectivity measures in the strict statistical-dependence sense. We note this expansion in interpretation in the Discussion.
4. In a number of cases, latent-factor operationalizations combining area-, household-, and individual-level indicators were used. These studies were retained if a census-derived indicator or an eligible area-level exposure was embedded in the latent factor. These studies were retained on the rationale that they represent a substantial and methodologically distinct subset of the literature. We acknowledge that the area-level contribution cannot always be cleanly isolated in these instances, and we discuss this caveat in the Discussion. Aggregate findings drawn from these studies should be interpreted with this constraint in mind.

## Results

The systematic database search yielded 3,241 records across PubMed (n=683), Embase (n=911), Scopus (n=1,082), and Web of Science (n=565), plus 2 additional records identified through Google Scholar, for a total of 3,243 records. After de-duplication in Covidence, 1,659 unique records were screened at the title and abstract level, of which 1,575 were excluded as ineligible if they were non-empirical (e.g., reviews, commentaries), did not report a validated area-level exposure or complementary place-based measure, or used a non-fMRI neuroimaging modality. Full-text review was conducted for 84 articles, resulting in the exclusion of 56 for the following reasons: studies using task-based fMRI only (n=26); non-empirical publications including preprints, perspectives, and reviews (n=7); use of a non-fMRI neuroimaging modality (n=6; including diffusion MRI [n=1], fNIRS [n=1], and structural MRI only [n=4]); area-level factors not assessed, unclear, or operationalized only at the individual or household level (n=14); no fMRI measure reported (n=1); duplicate record identified at full-text stage (n=1); and record withdrawn from the journal (n=1). A total of 28 studies met all inclusion criteria and were included in this review. The full PRISMA-ScR flow diagram is reported in **Figure 1**.

Included studies were published between 2017 and 2025. Seven studies were published in 2025, four in 2024, and three each in 2023 and 2022, and six in 2021 (**Figure 2B**). Fewer than one-third of included studies (k = 5, 17.9%) appeared in or before 2020, indicating that the field of understanding area-level neighborhood factors in neuroscience is growing in recent years ^50^. Coded by data origin, all 28 studies analyzed participants residing in the United States, despite first-author affiliations spanning five countries (United States, United Kingdom, Australia, Norway, and China). This pattern, in some part, reflects the field’s use of US data infrastructure, primarily the ABCD Study, and underscores the geographic concentration of the evidence base. Further, while the search was conducted across the full lifespan with no age restrictions, the available literature concentrates heavily in developmental samples (89.3% birth-through-18); we return to this in the Discussion as a finding of the review rather than a feature of its scope.

**Figure 2.**
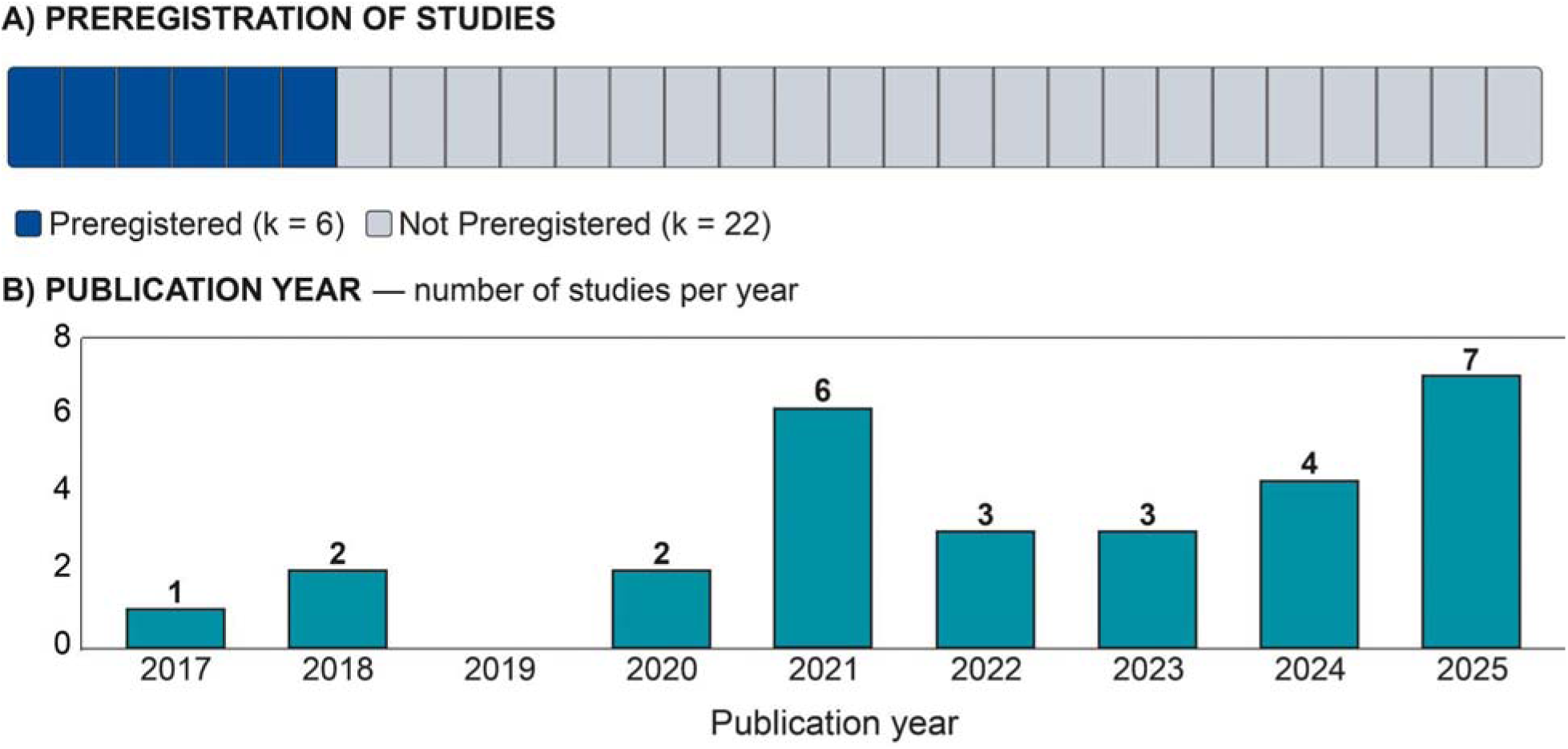
Preregistration of Included Studies and Publications Over Time. (A) Preregistration status of included studies (k = 28). The horizontal bar is divided into 28 equal segments, one per study; the 6 shaded segments correspond to studies with a publicly available pre-analysis registration (21.4%). (B) Number of studies identified by publication year.

The majority of included studies focused on developmental samples, leaving substantial gaps across the lifespan. Five studies examined perinatal or early-childhood populations; one study assessed fetal MRI ^51^, two studies assessed neonates cross-sectionally ^52,53^, and two studies used longitudinal designs extending from neonatal scans through toddlerhood or early childhood ^54,55^. The largest concentration of work focused on late childhood and early adolescence in samples drawn from the ABCD Study, accounting for 15 studies (53.6%). Of these, nine analyzed ABCD baseline data cross-sectionally at ages 9-10 ^49,56–63^, five examined baseline together with one or more follow-up waves ^64–68^; and one focused primarily on the year-2 follow-up (ages 11-12)^69^. Three studies spanned a wider range encompassing childhood-through-adolescence (approximately ages 6-19); one from the Michigan Twin Neurogenetics Study (MTwiNS) sample ^40^ and two single-site Detroit cohorts ^70,71^. Two studies focused on early adolescence specifically (ages 12-14; ^72,73^. Two studies extended into young adulthood (ages 5-25; ^74^, and ages 8-23 in the Philadelphia Neurodevelopmental Cohort (PNC) ^75^. Only one included study examined adults exclusively, drawing from a single-site post-trauma sample aged 18-65 ^76^. No included study focused on middle-aged or older adult populations. Taken together, 25 of the 28 included studies (89.3%) examined samples spanning the perinatal, childhood, or adolescent periods, with only three studies ^74–76^ extending meaningfully into adulthood and only one ^76^ examining adults exclusively. This concentration leaves substantial gaps across the lifespan.

Among studies that applied a parcellation scheme, the Gordon (333-region) atlas was by far the most commonly used (k = 15, 53.6%). Notably, this concentration likely reflects ABCD’s tabulated resting-state connectivity is released using Gordon-defined networks. The Tian subcortical atlas was applied in 4 studies (14.3%). Cerebellar atlases were used in 1 study (Kardan et al., 2025). The remaining 12 unique parcellation schemes (see ref: ^77^ for how these schemes differ in derivation and resolution) include the Harvard-Oxford, Shirer, Shen, AAL, and HCP-MMP atlases, as well as data-driven approaches (NMF, spectral clustering, InfoMap). This heterogeneity in parcellation schemes among 28 studies presents a challenge for cross-study synthesis, and raises questions regarding the sensitivity of results to the specific choice of parcellation. The primary direction and magnitude of associations between area-level factors and functional connectivity and their implications are discussed in detail below.

Lastly, individual study sample sizes ranged from 68 to 9,475 participants, with a median of 1,824 participants, reflecting marked variability driven largely by the inclusion of large epidemiological datasets (**Table 5**). The ABCD Study was by far the most commonly utilized data source (53.6%), establishing it as the dominant infrastructure for neighborhood functional neuroscience research in the United States. Independent or single-site cohorts accounted for 8 of the 28 studies (k = 8). The remaining studies drew from the Early Life Adversity and Biological Embedding (eLABE) study (k = 3) ^78^, the Philadelphia Neurodevelopmental Cohort (PNC; k = 1 ^79^), the Michigan Twin Neurogenetics Study (MTwiNS), and a neuroimaging follow-up to the Twin Study of Behavioral and Emotional Development (TBED-C; k = 1 ^80,81^).

Consistent with the broad eligibility criteria of this review, included studies employed a heterogeneous array of area-level measures, with some studies using multiple measures; see **Table 2** for full details. We organize this heterogeneity along two axes, (1) the substantive construct assessed (e.g., deprivation, crime, violence, opportunity) and (2) the geographic spatial unit used to capture it (e.g., geographic administrative index). Area-level factors were most commonly indexed by the Area Deprivation Index (ADI; k = 16 studies, 57.1%), including latent factors, making it the most frequently used validated composite deprivation measure. However, these 16 studies did not use the ADI in the same way. In 12, the ADI (or an ADI subcomponent) was modeled as a standalone, geocoded area-level predictor. In the remaining four, the ADI was embedded within a latent or multivariate composite that also combined household- and individual-level indicators (Beck et al., 2025; Modabbernia et al., 2021; Sripada et al., 2022; Tooley et al., 2024), such that its independent contribution could not be isolated. However, in the one study that tested the ADI separately from its composite, it showed no unique association with functional connectivity (Sripada et al., 2022). We therefore distinguish these two operationalizations when interpreting the direction of associations reported in **Table 3**.

**Table 2.** Characteristics of included studies (k = 28). Study-level summary of the 28 included studies, reporting first-author country of affiliation, the geographic unit of analysis at which area-level context was operationalized, and the specific area-level neighborhood measure(s) used. *Abbreviations:* UCR, Uniform Crime Reports

| Source | 1st-Author Affiliation | Unit of analysis | Area-Level Measure |
| --- | --- | --- | --- |
| Acosta-Rodriguez et al. 2025 | United States | Census tract | Area Deprivation Index |
| Beck et al. 2025 | Norway | Census tract | Latent factor including Area Deprivation Index |
| Brady et al. 2022 | United States | Census block group | UCR: Crime |
| Brady et al. 2024 | United States | Census block group | UCR: Crime |
| Cook et al. 2024 | United States | Census tract | Social Vulnerability Index |
| Ellwood-Lowe et al. 2021 | United States | County | UCR: Total violent crimes |
| Harris et al. 2023 | United States | Census tract | Child Opportunity Index |
| Iadipaolo et al. 2017 | United States | ZIP code | Distressed Communities Index |
| Ip et al. 2022 | United States | Census tract | Area Deprivation Index |
| Kardan et al. 2025 | United States | Census tract & County | Area Deprivation Index; UCR: Crime |
| Marshall et al. 2018 | United States | ZIP code | Distressed Communities Index |
| Michael et al. 2023 | United States | Census tract | Neighborhood poverty (tract level) |
| Michael et al. 2025 | United States | Census tract | Area Deprivation Index |
| Miller et al. 2018 | United States | Census block group | Neighborhood Murder Index |
| Miller et al. 2021 | United States | Census block group | Neighborhood Murder Index |
| Modabbernia et al. 2021 | United States | Census tract & County | Latent factor (CCA) including Area Deprivation Index; UCR: Crime |
| Rakesh et al. 2021a | Australia | Census tract | Area Deprivation Index |
| Rakesh et al. 2021b | Australia | Census tract | Area Deprivation Index |
| Rakesh et al. 2025a | United Kingdom | Census tract | Area Deprivation Index |
| Rakesh et al. 2025b | United Kingdom | State | Gini coefficient |
| Ramphal et al. 2020a | United States | Census block group | Area Deprivation Index |
| Ramphal et al. 2020b | United States | Census block group | Area Deprivation Index |
| Sripada et al. 2022 | United States | Census tract | Latent factor including Area Deprivation Index |
| Sydnor et al. 2023 | United States | Census block group | Latent factor using census crime data |
| Tooley et al. 2024 | United States | Census block group | Latent factor including Area Deprivation Index |
| Vargas et al. 2025 | United States | Census tract | Area Deprivation Index |
| Webb et al. 2021 | United States | Census block group | Area Deprivation Index |
| Zhao et al. 2024 | China | Census tract | Area Deprivation Index |

**Table 3.**
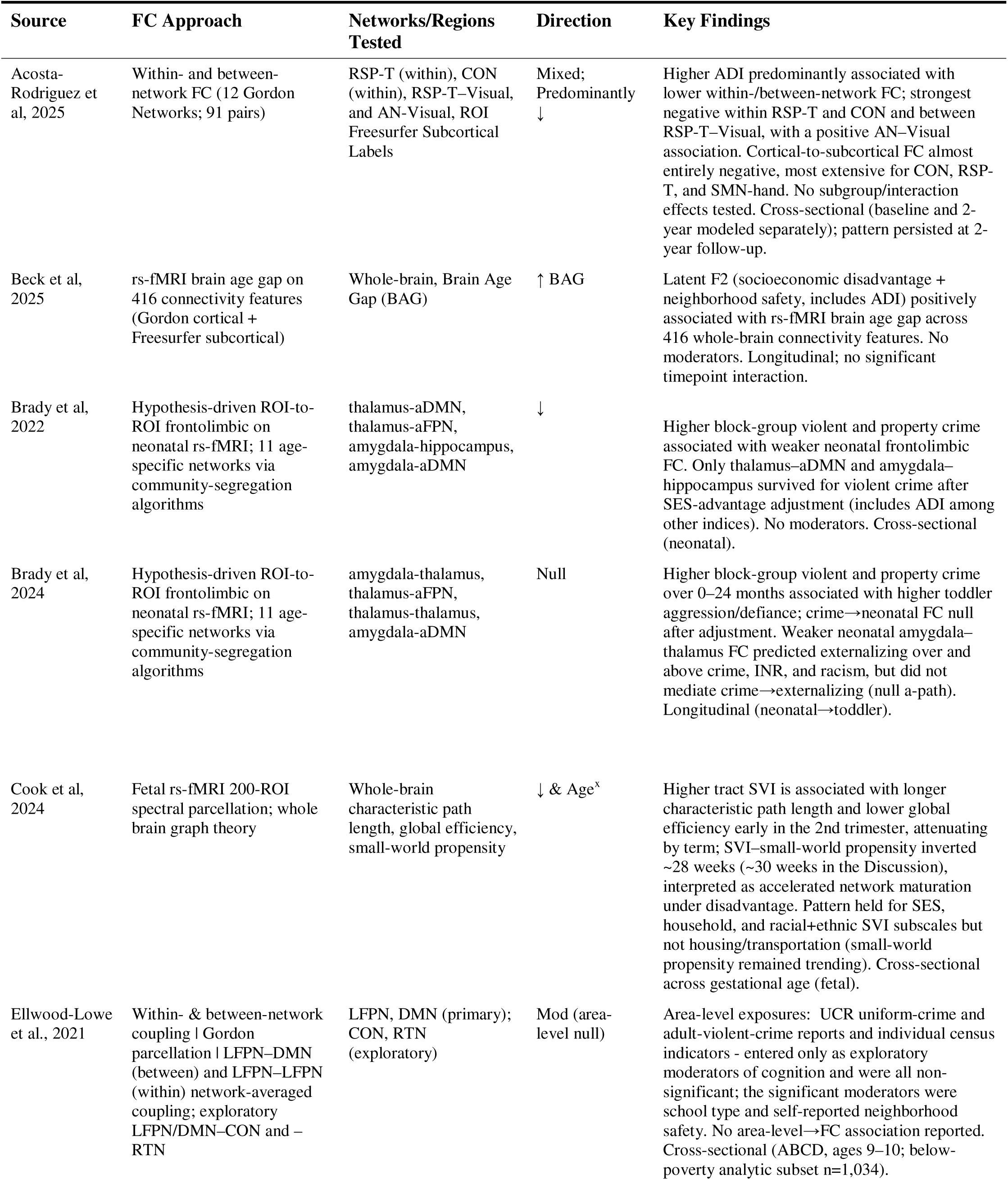
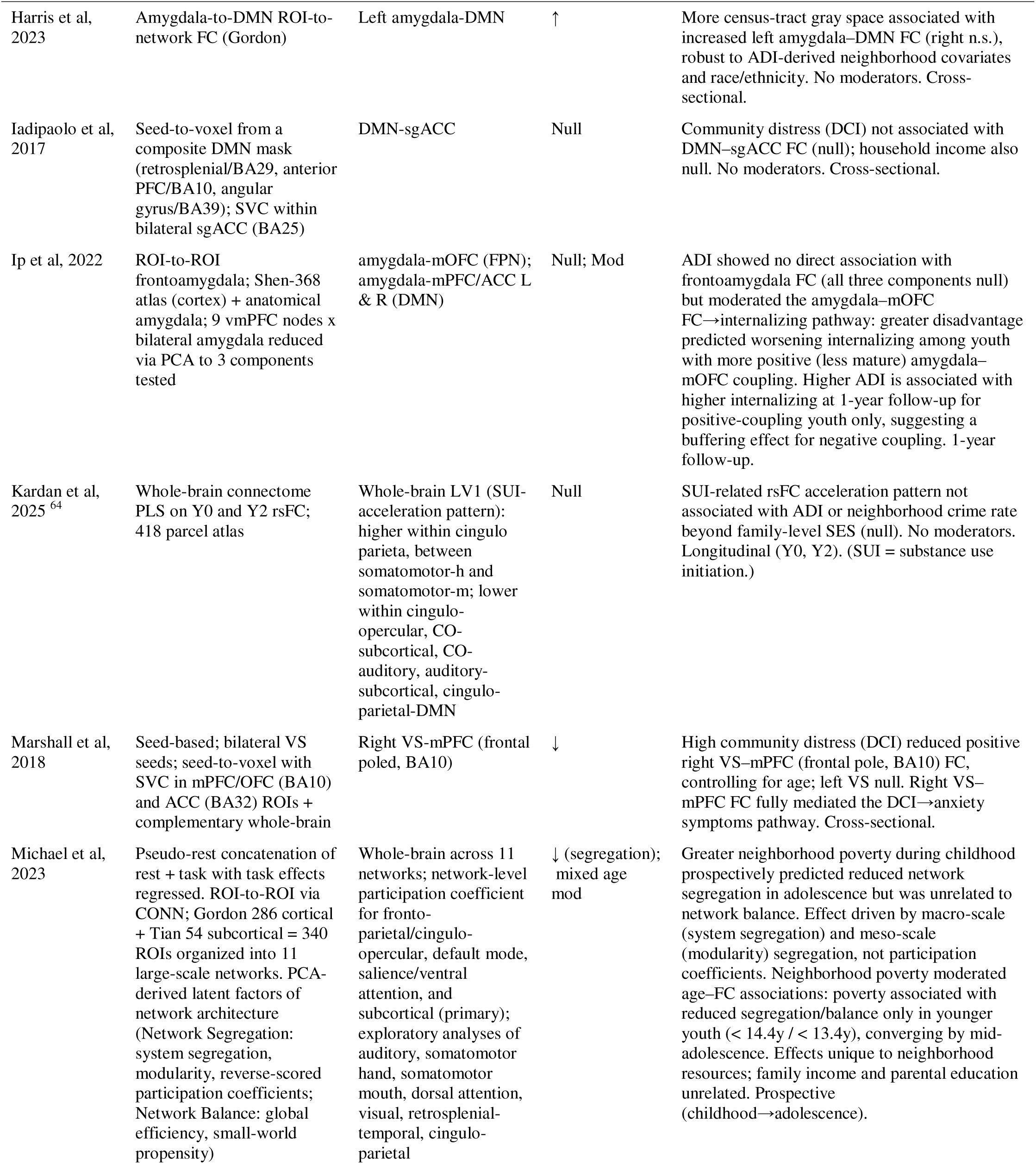
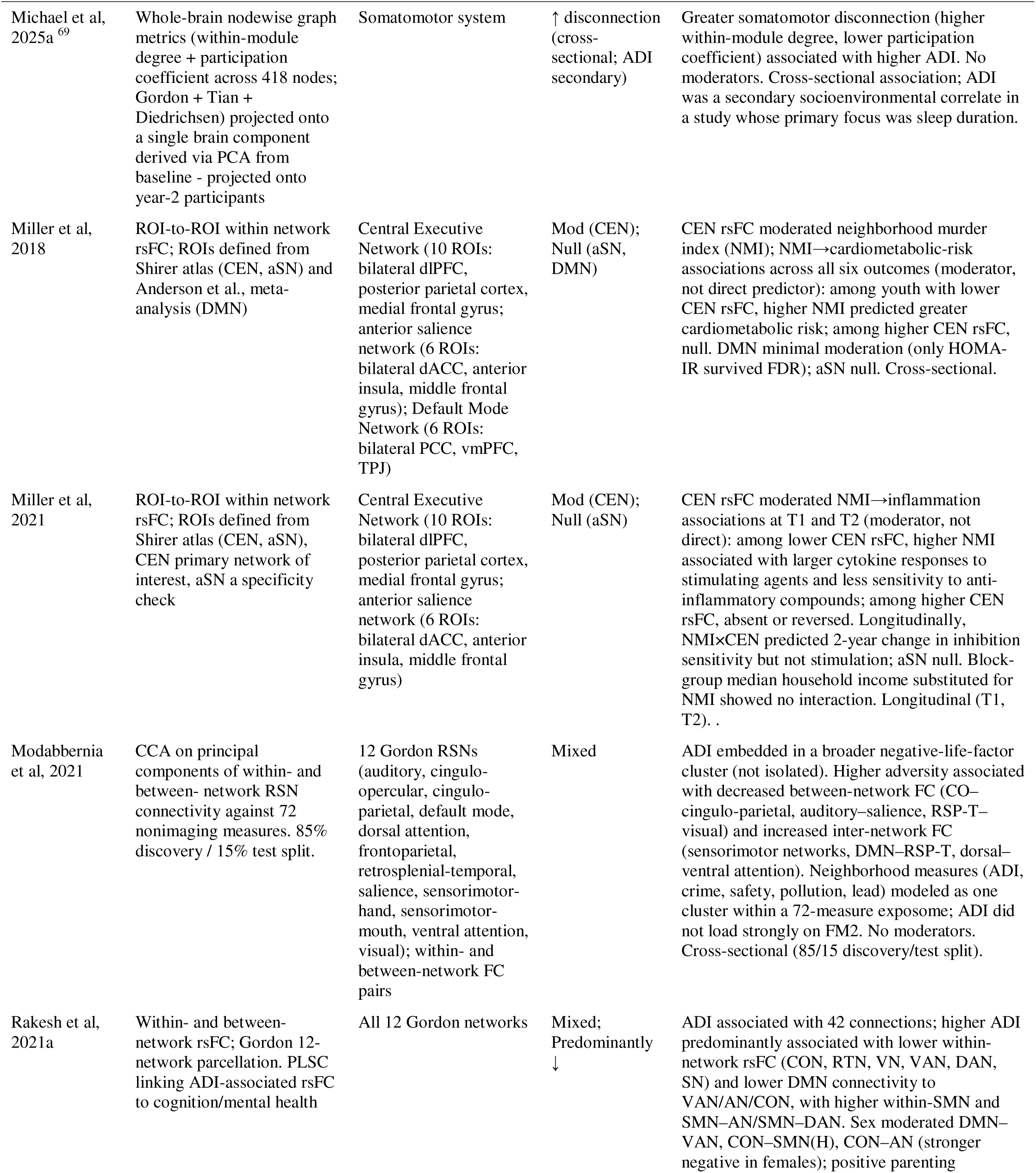
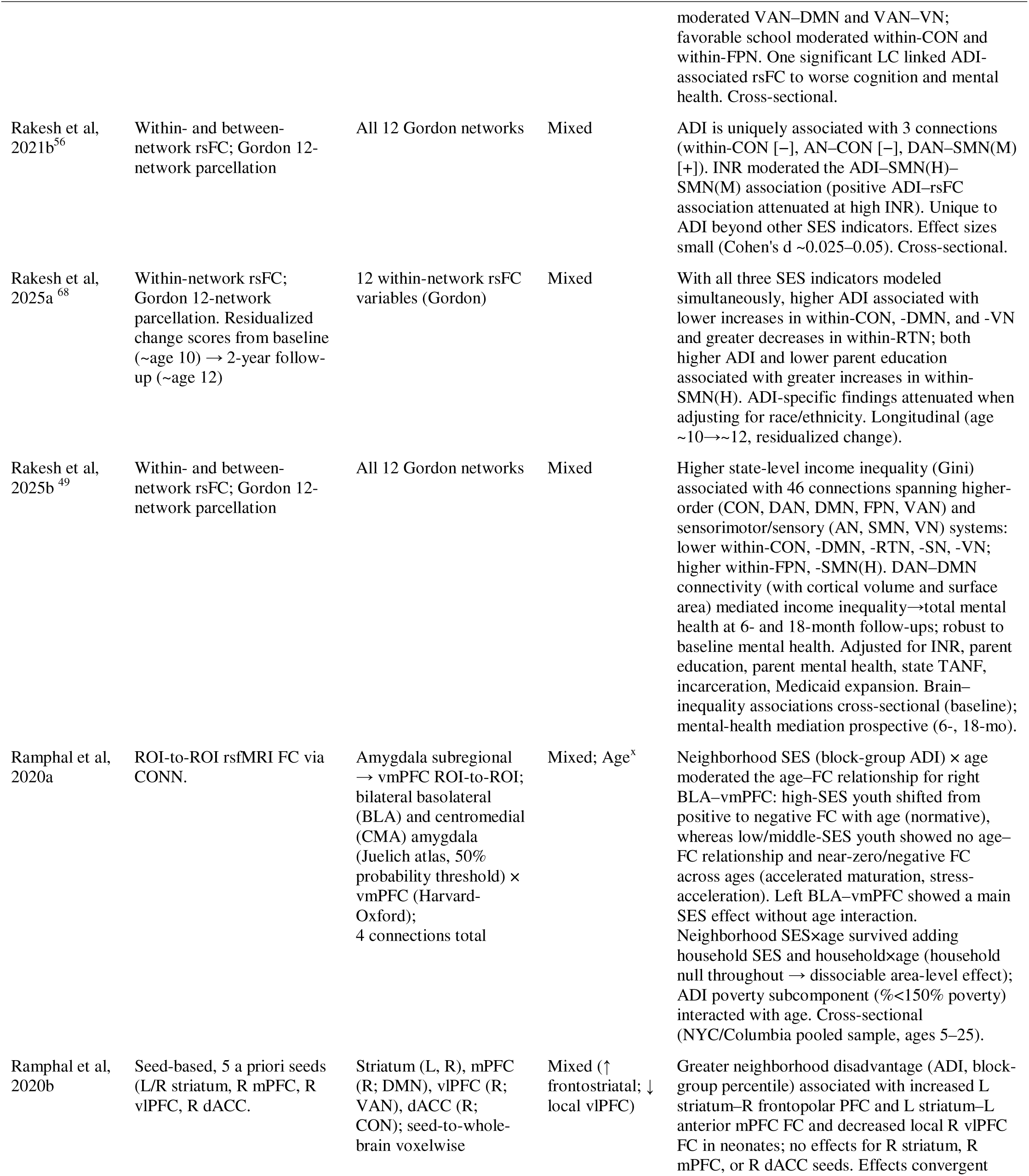
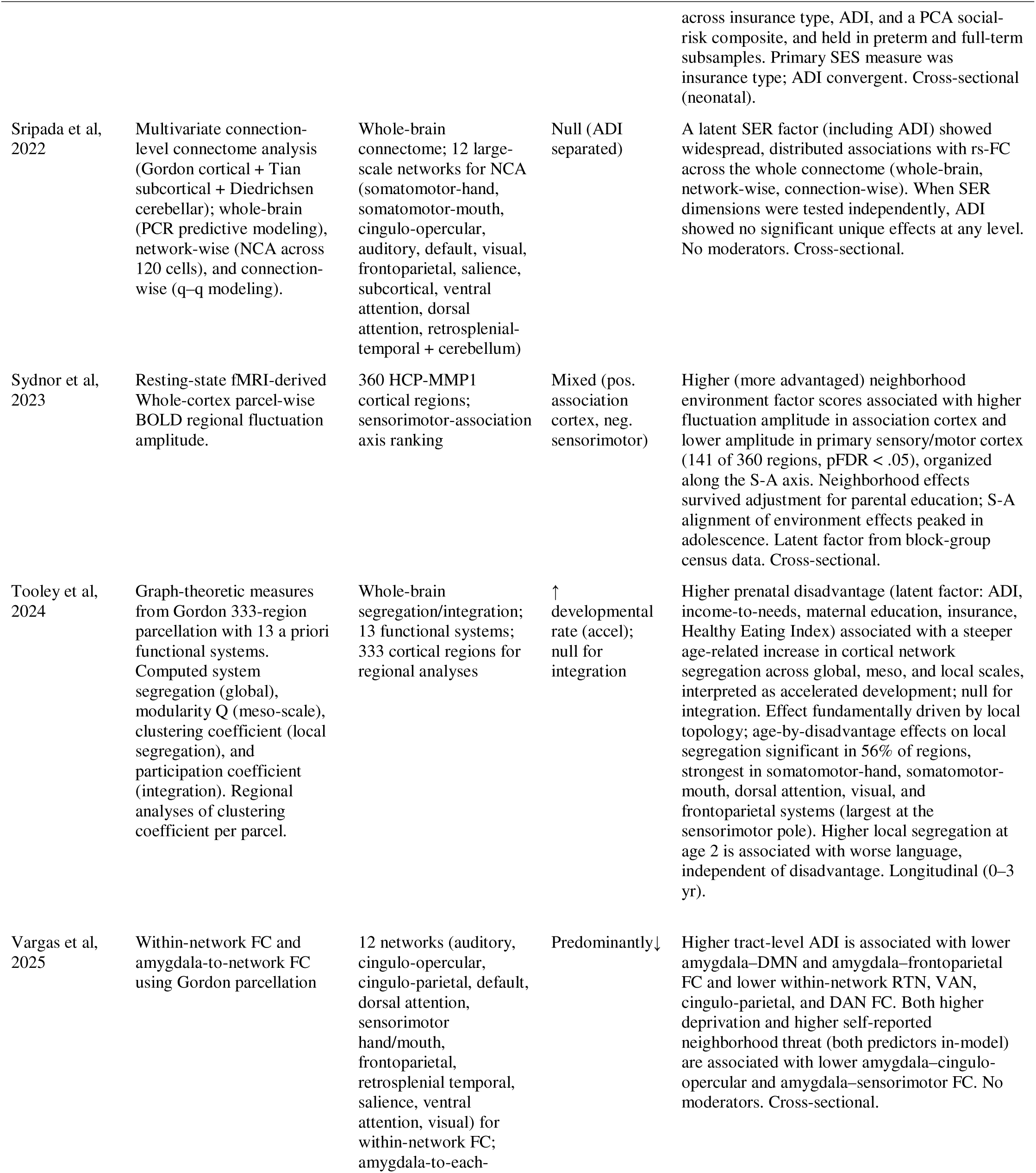
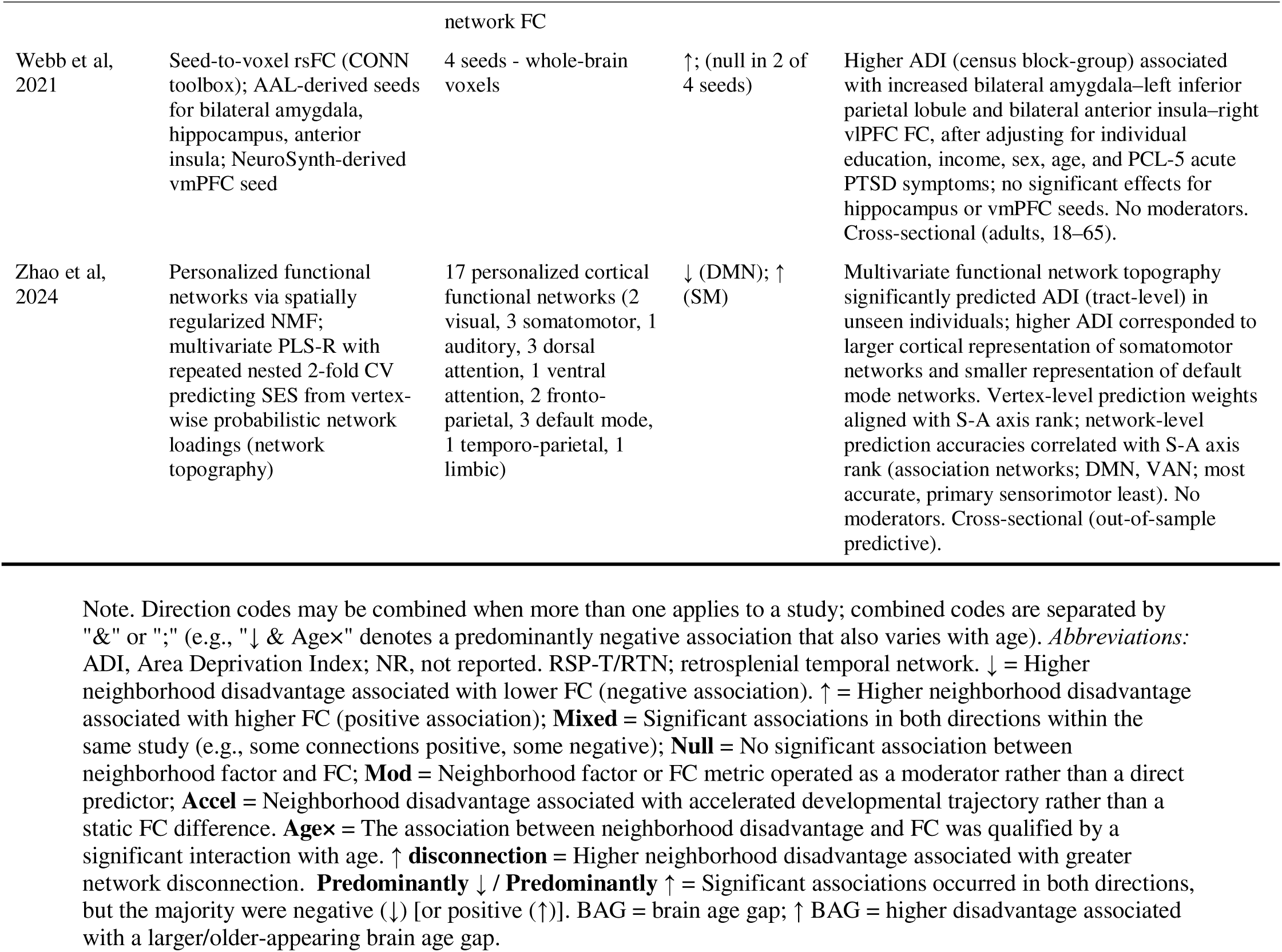
Functional connectivity findings of included studies (k = 28). Summary of the resting-state functional connectivity approach, the networks or regions tested, the coded direction of association with area-level disadvantage, and the key findings for each included study.

Less commonly used measures included the Distressed Communities Index (DCI; k = 2, 7.1%), the Social Vulnerability Index (SVI; k = 1, 3.6%), the Child Opportunity Index (COI; k = 1, 3.6%), and the Gini coefficient (k = 1, 3.6%). The Gini coefficient was not specified in the original preregistration but was captured here because it appeared alongside eligible area-level exposures. Geocoded exposure to area-level crime or violence was assessed in seven studies (25%) using a heterogeneous set of measures, including the FBI Uniform Crime Reporting Program ^48^, municipal police homicide records geocoded to participants’ addresses (e.g., Neighborhood Murder Index) ^72,73^, and the Applied Geographic Solutions CrimeRisk index ^82,83^, three of which combined crime exposure with the ADI ^54,63,64^. An additional set of studies (k = 6; see Supplementary Table S1) examined functional connectivity in relation to self- or informant-reported neighborhood-level constructs (e.g., Exposure to Violence questionnaire, Survey of Exposure to Community Violence, VEX-R) rather than geocoded administrative indices. Because these studies index respondents’ subjective experiences of community level exposures, rather than place-based conditions captured by the validated area-level indices that reflect the eligibility of this review ^29^, they did not meet the criteria for inclusion in the main synthesis. We catalog them in Supplementary Table S1 to document the broader landscape and to support future work explicitly comparing self-reported and geo-coded operationalizations of area-level vulnerability.

The geographic unit at which area-level context was operationalized varied substantially across studies (Figure 3). The most common resolution unit was the census tract used in 15 studies (53.6%), followed by the census block group used in 9 studies (32.1%). A smaller number of studies used ZIP code-level data (k = 2, 7.1%), county (k = 3, 10.7%), or state-level aggregates (k = 1, 3.6%). Notably, no study compared results across different levels of geographic resolution. Empirically, this variation is important given that census block groups (population ∼600–3,000) capture finer-grained residential contexts than census tracts (∼1,200–8,000) or ZIP codes (∼7,000–40,000), or counties (tens of thousands to millions), and the sensitivity of any given composite index to area-level effects may depend critically on the spatial grain at which area-level deprivation is measured.

**Figure 3:**
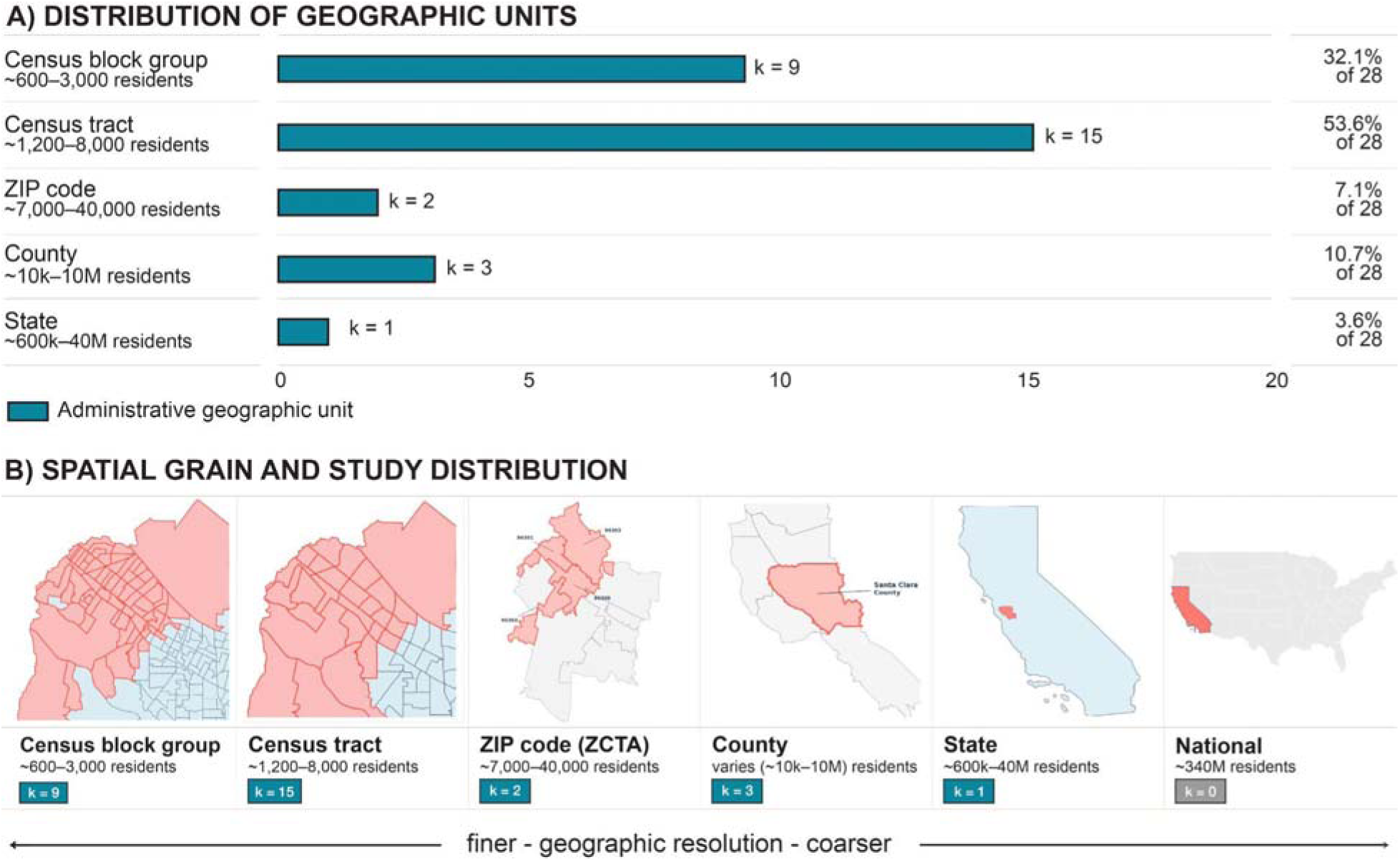
Distribution of Geographic Resolution Across Studies. Resolution of area-level measures across included studies (k = 28). (A) Number of studies using each spatial unit with percentages on the included set. (B) Spatial grain and study distribution (first author, year) for each geographic resolution; maps are illustrative and depict Palo Alto, California, United States, to show how a single location is partitioned at each spatial scale. Finer-grained units (block-group) reflect the smallest geographic unit, while coarser units (ZIP code, county, state, national) aggregate conditions across areas. Bars count each instance in which a resolution was used, so the two studies that used more than one resolution (Kardan et al., 2025; Modabbernia et al., 2021) contribute to more than one row; because these categories are not mutually exclusive, counts across rows do not sum to 100%.

### Direction and pattern of area-level associations with functional organization

Although the heterogeneity of measures, outcomes, and analytic approaches precludes meta-analytic pooling, several convergent patterns emerged across studies (Table 3). Over half of the corpus draws on overlapping ABCD baseline data most linked to the same area-level index (ADI), as such, recurrence across studies does not constitute independent replication.

The most consistent direction of effect was a negative association between higher area-level disadvantage and within-network connectivity in higher-order association and control systems. Higher ADI was associated with lower within-network connectivity in cingulo-opercular, default mode, retrosplenial-temporal, visual, ventral attention, and dorsal attention networks across multiple ABCD analyses ^56,61,65^, a pattern echoed for state-level income inequality ^49^. Running counter to this, somatomotor connectivity was frequently elevated under greater disadvantage (higher within-somatomotor connectivity in Rakesh et al., 2021a^57^, 2021b^56^, 2025a^68^; greater somatomotor ‘disconnection’ with higher ADI in Michael et al., 2025^69^; and larger cortical representation of somatomotor networks with higher ADI in Zhao et al., 2024^58^). This association/control-down, somatomotor-up dissociation was a within-ABCD, within-ADI pattern. However, because these analyses are almost entirely ABCD-based and ADI-indexed, this convergence should be read as a within-dataset, within-instrument pattern.

Three studies independently reported that area-level effects on intrinsic organization are patterned along the sensorimotor-to-association axis. Neighborhood-environment factor scores were associated with regional BOLD fluctuation amplitude in a manner organized along the S-A axis, with alignment peaking in adolescence ^75^; multivariate prediction weights for ADI aligned with S-A axis rank, with association networks most predictive ^58^; and age-by-disadvantage effects on local segregation were largest at the sensorimotor pole ^55^.

Findings for amygdala- and striatal-based circuits were more mixed and frequently moderation-based. Higher disadvantage was associated with lower amygdala–default mode and amygdala-frontoparietal connectivity in adolescence ^61^ and weaker neonatal frontolimbic connectivity following higher block-group crime exposure ^53^, whereas other studies reported higher amygdala-based connectivity ^76^ or effects contingent on protective features of the environment ^62^. Several frontolimbic findings were interpreted through an accelerated-maturation lens (see below) rather than as static differences ^52,74^. A recurring developmental interpretation, rather than a uniform main effect, was that area-level disadvantage is associated with an accelerated or compressed maturational trajectory. This framing appeared for corticolimbic connectivity ^74^, fetal network maturation ^51^, and the age-related increase in cortical network segregation in early childhood ^55^, and is consistent with reports that poverty-related reductions in network segregation were present in younger but not older adolescents, converging by mid-adolescence ^40^. These studies span the perinatal-through-adolescent range, but direct cross-period comparisons within a single design remain absent. Community distress was unrelated to default-mode-subgenual connectivity ^70^, and ADI showed no direct association with frontoamygdala connectivity while still moderating an internalizing pathway ^67^. In two studies, central-executive-network connectivity moderated associations between neighborhood crime and cardiometabolic or inflammatory outcomes without a direct connectivity effect ^72,73^.

Critically, several headline associations attenuated or disappeared once area-level exposure was statistically isolated from household-level socioeconomic status; ADI showed no unique connectome-wide effect when separated from a broader socioeconomic-risk latent factor ^59^, and an acceleration pattern was not associated with ADI beyond family-level SES ^64^. Findings from ADI-specific analyses also attenuated after adjustment for race/ethnicity ^68^. A substantial subset of studies, however, reported null direct associations or effects that operated only through moderation (e.g., ^59,60,67,70,72,73^, underscoring the difficulty of disentangling area-level signal from confounding individual and household conditions.

Lastly, a smaller set of studies using graph-theoretic metrics converged in suggesting that disadvantage is associated with altered global and local topology; longer characteristic path length and lower global efficiency early in gestation ^51^, steeper age-related increases in segregation ^55^, and reduced network segregation in adolescence ^40^; though the scarcity and heterogeneity of these measures (Section 4) limits firm conclusions. Taken together, the literature is directionally suggestive, most consistently for a within-association-network reduction coupled with somatomotor elevation, an S-A-axis organization, and an accelerated-development motif but the dependence of these patterns on a shared dataset and instrument, the prevalence of null or moderation-only direct effects, and the attenuation of effects once household SES and race/ethnicity are accounted for, all caution against strong mechanistic or generalizable conclusions at this stage.

## Discussion

This scoping review set out to map the empirical literature on associations between area-level factors and resting-state functional brain connectivity and network organization across the lifespan. Across 28 identified studies, we characterized the range of area-level exposures employed, the functional connectivity outcomes assessed, the populations studied, and the analytic approaches used. At the same time, the field is expanding rapidly, heavily concentrated in developmental samples, and methodologically heterogeneous in how area-level context and functional connectivity are operationalized. Below, we discuss these observations with attention to both their implications for the current state of the field and the opportunities they identify for future research.

### 1. Recurring Patterns in Reviewed Findings

Because this is a scoping review, we mapped rather than formally synthesized the literature: we did not appraise study quality or pool effect sizes, and over half of the corpus draws on overlapping ABCD data most linked to a single index. The observations below are therefore descriptive patterns across the reviewed studies, not weighted conclusions about consistency or magnitude. With that caveat, a few patterns recurred across otherwise distinct samples. Several studies reported lower connectivity within higher-order association and control networks, most often the cingulo-opercular and retrosplenial-temporal networks, alongside increased or atypical somatomotor connectivity under greater disadvantage ^56–58,61,65,69^. In several cases this patterning was described along the sensorimotor-to-association axis ^55,58,75^. A developmental framing also recurred, with disadvantage-related effects interpreted as accelerated maturation rather than static differences ^51,55,74^. Frontolimbic circuitry was the most frequently studied target but showed the least directionally consistent results, often appearing as age-dependent or moderating effects. These observations remain preliminary as reported effects were typically small, with several attenuated once area-level exposure was separated from household socioeconomic status or race/ethnicity, and apparent convergence partly reflects use of common data rather than independent replication underscoring the methodological priorities we turn to next.

### 2. Measurement Heterogeneity in Area-Level Exposures

One of the most striking features of this literature is the wide variation in how area-level factors are operationalized. Across included studies, neighborhood disadvantage was indexed using approaches ranging from census-derived deprivation composites to fully geocoded, validated multidimensional indices such as the Area Deprivation Index (ADI), Child Opportunity Index (COI), and Social Vulnerability Index (SVI); A subset of studies (k = 4; ^55,59,63,66^ combined ADI with individual- or household-level socioeconomic indicators (e.g., income, parental education) into latent factors. Although beyond the scope of this review, single-item self-report measures of perceived neighborhood safety or quality were commonly investigated in tandem with the area-level indices ^55,60,61,68^. This heterogeneity may reflect a conceptual ambiguity about what area-level context means across studies. Future work would benefit from reporting area-level effects both within and independent of latent contextual factors.

Self-report measures of perceived neighborhood quality capture a psychological construct of how residents experience and appraise their environment. That is distinct from the neighborhood, place-based conditions indexed by geocoded composite measures. Treating these interchangeably conflates mechanisms and limits the interpretability of findings. Moreover, even among studies using geocoded indices, substantial variation exists in the domains captured: deprivation-focused measures such as the ADI emphasize socioeconomic deficits, while opportunity-based measures such as the COI additionally capture access to educational, health, and environmental resources, a conceptual distinction that may be important for capturing the developmental contexts most relevant to neural and cognitive outcomes. Moving forward, the growing field would benefit considerably from greater theoretical intentionality and standardization in the selection of area-level measures, guided by the constructs and mechanisms under investigation.

Beyond improvements in measurement, the rigor of inferences in this literature would be strengthened by greater uptake of preregistration. Among the 28 studies included in this review, six reported a preregistered analysis plan^55–57,60,61,68^ (**Figure 2A**). Preregistration of hypotheses, exposure operationalizations, and pre-planned sensitivity analyses (including across geographic resolutions) would improve the replicability of reported associations between area-level context and functional brain organization. When confirmatory designs are not feasible, transparent acknowledgment of exploratory framing is a step toward methodological clarity.

### 3. Geographic Resolution as a Distinct Dimension of Measurement Heterogeneity

Beyond the choice of which area-level index is used, another dimension of measurement heterogeneity is the geographic resolution at which area-level context is operationalized. Across included studies, the same constructs - area deprivation, crime exposure, socioeconomic opportunity - were measured at spatial scales ranging from the census block group (∼600–3,000 residents) to the census tract (∼1,200–8,000), ZIP code (∼7,000–40,000), and in some cases the county or state level (**Figure 3**). The block group and census tract capture meaningfully different aspects of residential context, with finer-grained units reflecting the immediate street-level environment most proximal to daily lived experience, while coarser units aggregate conditions across heterogeneous areas that may dilute or obscure within-neighborhood variation ^84,85^. Critically, no study in this review explicitly justified its choice of geographic unit, and none tested whether results were robust to resolution, leaving open the question of whether reported associations are robust features of the exposure-brain relationship or partly artifacts of the spatial scale at which area-level context was operationalized. This is a version of the modifiable areal unit problem (MAUP; ^84^), a well-documented challenge in spatial epidemiology, comprising both scale and zonation effects, that has received less attention in the neuroimaging literature. Scale effects, in which results change as units are aggregated from finer to coarser resolutions (e.g., block group to tract to county), compound with zonation effects, in which results change as boundaries are redrawn at the same resolution. Empirical work in spatial epidemiology has shown the magnitude and even the direction of area-level health associations can shift with the geographic unit selected, and therefore the appropriate unit is the one matched to the hypothesized causal process and spatial structure of the exposure ^86–88^.

A tractable methodological response is the adoption of multiverse analysis, systematically re-estimating models across all plausible combinations of area-level index and geographic resolution, and reporting the full distribution of resulting effect sizes ^89,90^. Applied to area-level factors in neuroscience, a multiverse approach would reveal whether associations between neighborhood context and functional connectivity are robust to measurement decisions or are sensitive to the particular index-resolution combination chosen, a finding with direct implications for both replication and policy translation. We recommend that future studies report at minimum a sensitivity analysis testing whether effects replicate at an adjacent level of geographic aggregation, and provide an explicit justification of the spatial unit(s) of analysis in studies of neighborhood context and brain outcomes.

Our eligibility criteria were scoped to the Trinidad et al. (2022) inventory of area-level deprivation measures used in US public health research, supplemented by the Opportunity Atlas ^46,47^ and UCR-based crime exposure ^48^. By design, this scope does not capture area-level deprivation instruments developed and commonly used outside the United States context. The US-centric measurement infrastructure reviewed here reflects the scope of the Trinidad et al. 2022 inventory on which the eligibility criteria of this scoping review were based. Further work cataloging non-US area-level deprivation measures, including but not limited to, the Canadian Index of Multiple Deprivation and Québec-specific Material and Social Deprivation Index (MSDI) ^91,92^, New Zealand NZDep2013 Index of Deprivation ^93^, Australia’s Socio-Economic Indexes for Areas (SEIFA) ^94,95^, European and pan-European ^96–98^, Japanese Areal Deprivation Index ^99^, and their uptake in neuroimaging would be a valuable component in future work. Further, cross-national comparison, e.g., across contexts differing in welfare provision and/or housing systems, would test whether these associations reflect a generalizable construct or features specific to the US, and would motivate harmonizing area-level exposure measures across international neuroimaging consortia.

### 4. Underutilization of Whole Brain Network Metrics and Analytic Approaches

A related methodological gap concerns the level at which functional brain organization is characterized. The majority of studies identified in this review examined functional connectivity at the level of pairwise connections between regions of interest, or within- and between-network connectivity among predefined large-scale networks. While informative, these approaches capture local or network-specific relationships and may miss the global organizational properties of the brain especially given that the recurring premise of this literature is that chronic, diffuse exposures such as sustained deprivation act on the brain as a system rather than on isolated connections. Graph-theoretic metrics are one such family, e.g., modularity (the degree to which the network separates into distinct communities), the clustering coefficient (local segregation), and characteristic path length or global efficiency (integration, or how readily information traverses the network) ^100,101^. These metrics are particularly well-suited to detecting how conditions that operate at the level of whole systems, such as sustained deprivation or chronic threat exposure, may reorganize the brain’s large-scale architecture. For example, reduced network modularity, reflecting less distinct separation between functional systems, has been associated with cognitive difficulties and adversity exposure in prior work ^102,103^, yet this class of measure remains largely absent from the area-level literature. Other approaches are similarly underrepresented, such as time-varying connectivity and state-space models that capture how connectivity reconfigures over time, individualized parcellation and precision functional mapping. No single method is uniquely sensitive, and each carries its own dependencies on parcellation, thresholding, and motion; the broader point is that wider adoption of whole-network and temporally resolved approaches would more fully characterize how area-level context shapes not just the strength of specific connections but the global organization of functional brain architecture across development.

### 5. Homogeneity of Analytic Samples

The studies identified in this review draw on participant samples that are demographically homogeneous and inconsistently characterized (**Table 4**). The majority of included samples were predominantly non-Hispanic White, with this group exceeding 60% in most studies using ABCD. More racially and ethnically diverse samples were largely confined to single-site cohorts that recruited from urban contexts ^52,55,70–73,76^, a pattern that confounds sample demographics with statistical power, geographic specificity, and study design. This heterogeneity in reporting racial and ethnic sample composition may constrain synthesis across studies and the broader collective inferences made from this work. Further, this pattern may be compounded by analytic decisions that can occur after recruitment. For example, head motion correction and motion-based participant exclusion are necessary for valid functional connectivity estimation ^104–107^. However, cross-sectional work has shown that minoritized youth in ABCD exceed standard motion thresholds at higher rates than White youth at baseline ^106^, and recent longitudinal analyses extend this observation; Black, Hispanic, and Asian participants were more likely to be excluded for motion at baseline (ages 9-10), and Black participants were less likely to return for two-year follow-up after adjusting for socioeconomic factors and baseline motion ^108^. Because over half of the studies included in this review draw on ABCD, the populations to whom the field’s inferences apply are narrowed at upstream stages of the analytic pipeline. This narrowing has implications for the literature focused on area-level factors, where associations between deprivation indices and functional connectivity are estimated within samples increasingly unlike the communities in which area-level disadvantage is concentrated. Addressing this gap will require reporting quality-control and evaluating the sensitivity of findings to motion exclusion thresholds, and lastly considering how these analytic frameworks are designed to retain rather than discard high-motion participants where the scientific question warrants ^109^.

**Table 4.**
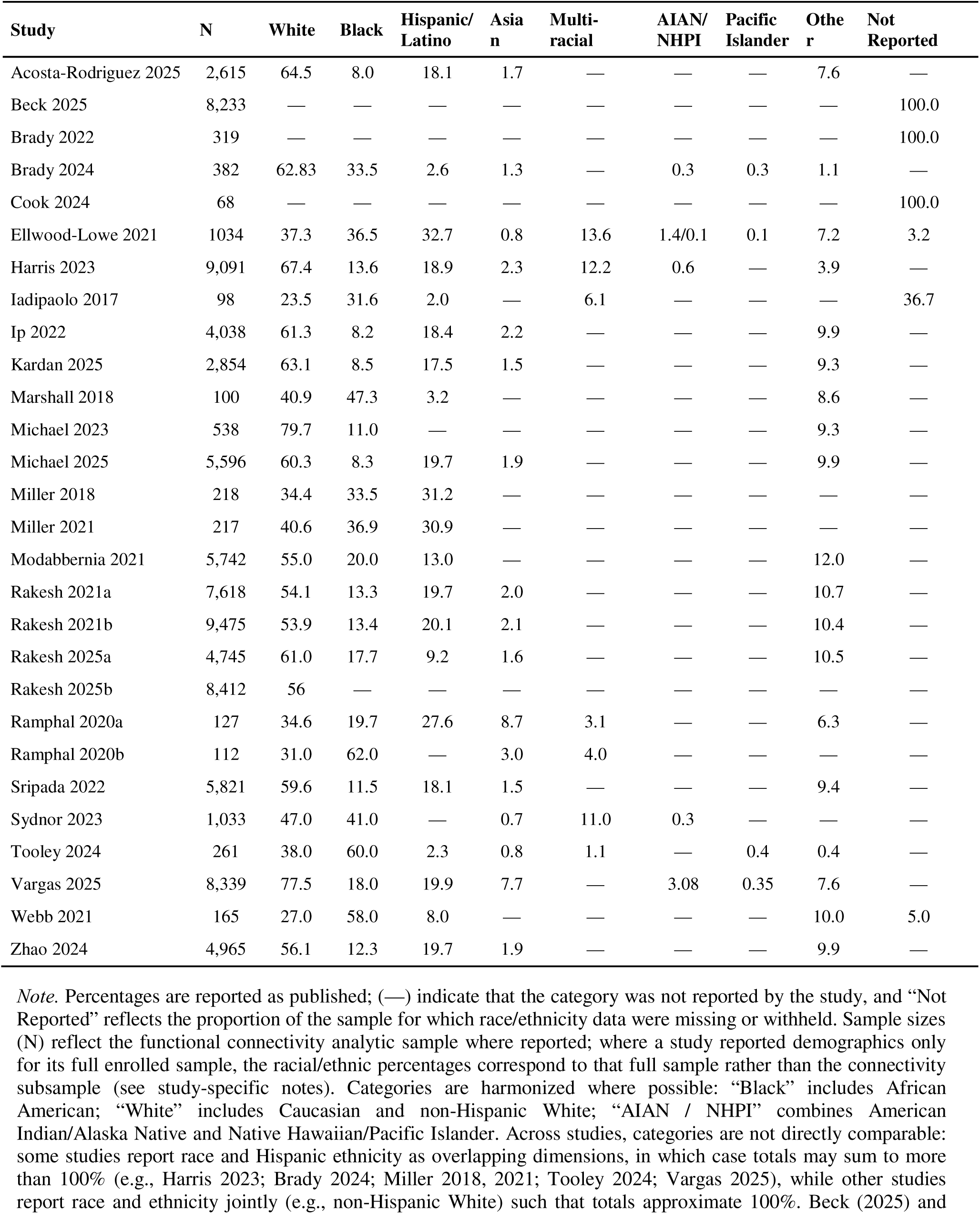
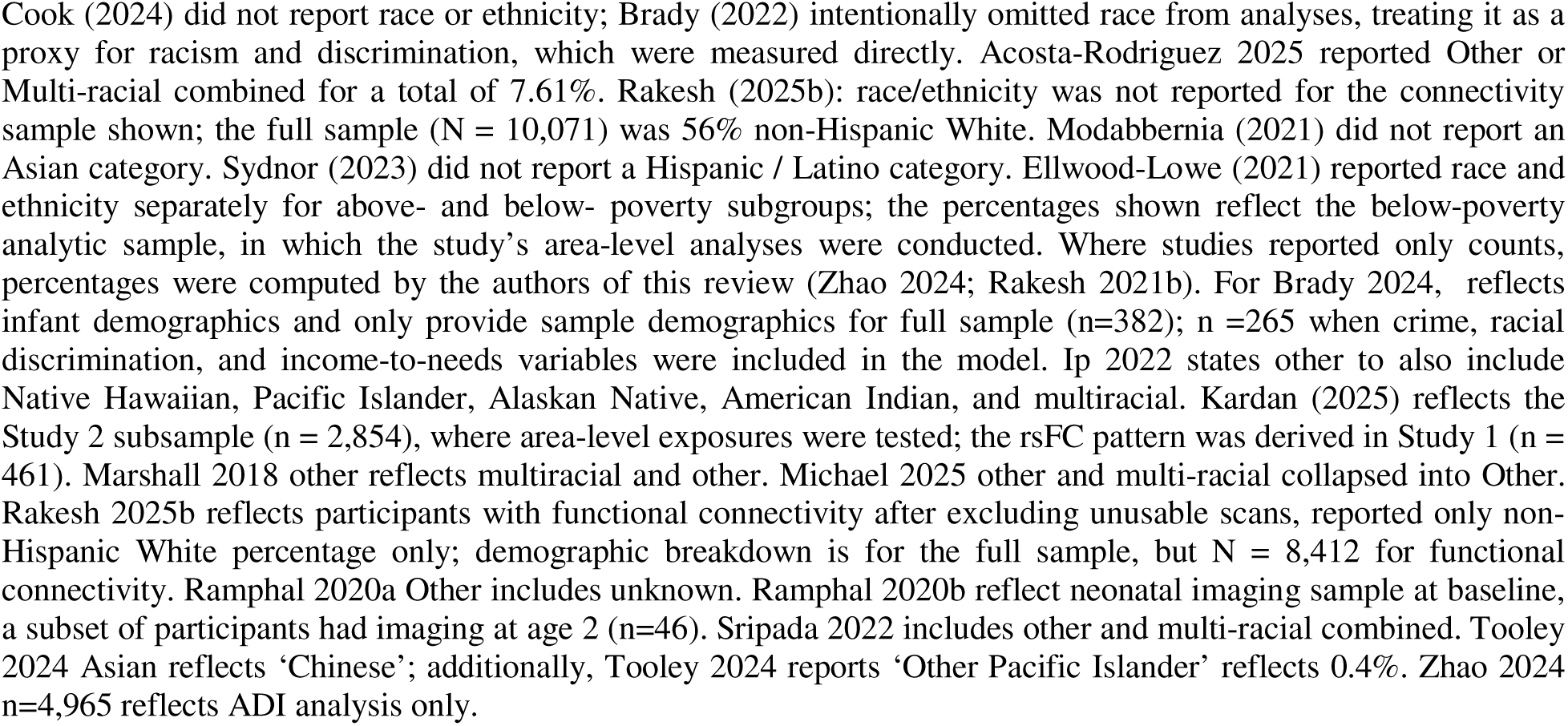
Racial and ethnic distribution of participants across included studies (%). Per-study analytic sample size and racial/ethnic composition, harmonized across studies where possible.

| Study | N | White | Black | Hispanic/<br>Latino | Asian | Multi-<br>racial | AIAN/<br>NHPI | Pacific<br>Islander | Other | Not<br>Reported |
| --- | --- | --- | --- | --- | --- | --- | --- | --- | --- | --- |
| Acosta-Rodriguez 2025 | 2,615 | 64.5 | 8.0 | 18.1 | 1.7 | — | — | — | 7.6 | — |
| Beck 2025 | 8,233 | — | — | — | — | — | — | — | — | 100.0 |
| Brady 2022 | 319 | — | — | — | — | — | — | — | — | 100.0 |
| Brady 2024 | 382 | 62.83 | 33.5 | 2.6 | 1.3 | — | 0.3 | 0.3 | 1.1 | — |
| Cook 2024 | 68 | — | — | — | — | — | — | — | — | 100.0 |
| Ellwood-Lowe 2021 | 1034 | 37.3 | 36.5 | 32.7 | 0.8 | 13.6 | 1.4/0.1 | 0.1 | 7.2 | 3.2 |
| Harris 2023 | 9,091 | 67.4 | 13.6 | 18.9 | 2.3 | 12.2 | 0.6 | — | 3.9 | — |
| Iadipaolo 2017 | 98 | 23.5 | 31.6 | 2.0 | — | 6.1 | — | — | — | 36.7 |
| Ip 2022 | 4,038 | 61.3 | 8.2 | 18.4 | 2.2 | — | — | — | 9.9 | — |
| Kardan 2025 | 2,854 | 63.1 | 8.5 | 17.5 | 1.5 | — | — | — | 9.3 | — |
| Marshall 2018 | 100 | 40.9 | 47.3 | 3.2 | — | — | — | — | 8.6 | — |
| Michael 2023 | 538 | 79.7 | 11.0 | — | — | — | — | — | 9.3 | — |
| Michael 2025 | 5,596 | 60.3 | 8.3 | 19.7 | 1.9 | — | — | — | 9.9 | — |
| Miller 2018 | 218 | 34.4 | 33.5 | 31.2 | — | — | — | — | — | — |
| Miller 2021 | 217 | 40.6 | 36.9 | 30.9 | — | — | — | — | — | — |
| Modabbernia 2021 | 5,742 | 55.0 | 20.0 | 13.0 | — | — | — | — | 12.0 | — |
| Rakesh 2021a | 7,618 | 54.1 | 13.3 | 19.7 | 2.0 | — | — | — | 10.7 | — |
| Rakesh 2021b | 9,475 | 53.9 | 13.4 | 20.1 | 2.1 | — | — | — | 10.4 | — |
| Rakesh 2025a | 4,745 | 61.0 | 17.7 | 9.2 | 1.6 | — | — | — | 10.5 | — |
| Rakesh 2025b | 8,412 | 56 | — | — | — | — | — | — | — | — |
| Ramphal 2020a | 127 | 34.6 | 19.7 | 27.6 | 8.7 | 3.1 | — | — | 6.3 | — |
| Ramphal 2020b | 112 | 31.0 | 62.0 | — | 3.0 | 4.0 | — | — | — | — |
| Sripada 2022 | 5,821 | 59.6 | 11.5 | 18.1 | 1.5 | — | — | — | 9.4 | — |
| Sydnor 2023 | 1,033 | 47.0 | 41.0 | — | 0.7 | 11.0 | 0.3 | — | — | — |
| Tooley 2024 | 261 | 38.0 | 60.0 | 2.3 | 0.8 | 1.1 | — | 0.4 | 0.4 | — |
| Vargas 2025 | 8,339 | 77.5 | 18.0 | 19.9 | 7.7 | — | 3.08 | 0.35 | 7.6 | — |
| Webb 2021 | 165 | 27.0 | 58.0 | 8.0 | — | — | — | — | 10.0 | 5.0 |
| Zhao 2024 | 4,965 | 56.1 | 12.3 | 19.7 | 1.9 | — | — | — | 9.9 | — |

*Note.* Percentages are reported as published; (—) indicate that the category was not reported by the study, and “Not Reported” reflects the proportion of the sample for which race/ethnicity data were missing or withheld. Sample sizes (N) reflect the functional connectivity analytic sample where reported; where a study reported demographics only for its full enrolled sample, the racial/ethnic percentages correspond to that full sample rather than the connectivity subsample (see study-specific notes). Categories are harmonized where possible: “Black” includes African American; “White” includes Caucasian and non-Hispanic White; “AIAN / NHPI” combines American Indian/Alaska Native and Native Hawaiian/Pacific Islander. Across studies, categories are not directly comparable: some studies report race and Hispanic ethnicity as overlapping dimensions, in which case totals may sum to more than 100% (e.g., Harris 2023; Brady 2024; Miller 2018, 2021; Tooley 2024; Vargas 2025), while other studies report race and ethnicity jointly (e.g., non-Hispanic White) such that totals approximate 100%. Beck (2025) and

**Table 5.** Datasets utilized across included studies (k = 28). Included studies grouped by underlying data source (ABCD, eLABE, MTwiNS, PNC, and independent/single-site cohorts), with a description of each dataset’s design, age range, and relevance to the review.

| Dataset | Studies | Description |
| --- | --- | --- |
| Adolescent Brain Cognitive Development (ABCD) | Acosta-Rodriguez 2025; Beck 2025; Ellwood-Lowe 2021; Harris 2023; Ip 2022; Kardan 2025; Michael 2025; Modabbernia 2021; Rakesh 2021a; Rakesh 2021b; Rakesh 2025a; Rakesh 2025b; Sripada 2022; Vargas 2025; Zhao 2024 | The largest long-term study of brain development and child health in the United States, enrolling approximately 11,800 children aged 9–10 at baseline across 21 sites with planned biennial follow-up into early adulthood. Its scale, geographic spread, and linkage of residential addresses to area-level indices (notably the ADI) have made it the dominant infrastructure for neighborhood functional neuroscience. |
| Early Life Adversity and Biological Embedding (eLABE) | Brady 2022; Brady 2024; Ramphal 2020b | A single-site longitudinal birth cohort (Washington University, St. Louis) oversampling for prenatal social disadvantage, with neonatal resting-state fMRI acquired shortly after birth and follow-up through toddlerhood and early childhood. Its perinatal focus makes it a primary source for the review’s small cluster of fetal/neonatal studies, examining how area-level exposures relate to frontolimbic connectivity early in development. |
| Michigan Twin Neurogenetics Study (MTwiNS) | Michael 2023 | A neuroimaging follow-up to the Twin Study of Behavioral and Emotional Development (TBED-C), recruiting twin pairs from Michigan birth records with oversampling from economically disadvantaged neighborhoods. Its twin design and intentional sampling across the poverty distribution distinguish it from convenience cohorts, and it spans a wide childhood-through-adolescence age range (~6–19). |
| Philadelphia Neurodevelopmental Cohort (PNC) | Sydnor 2023 | A single-site cross-sectional cohort of youth (~8–23 years) recruited through the Children’s Hospital of Philadelphia, with imaging and neighborhood-linked census/crime data. It contributes one of the few studies extending meaningfully into young adulthood and one of the few using a sensorimotor–association axis framing of intrinsic activity. |
| Independent / single-site cohorts | Cook 2024; Iadipaolo 2017; Marshall 2018; Miller 2018; Miller 2021; Ramphal 2020a; Tooley 2024; Webb 2021 | Eight studies drawing on individually recruited samples rather than large shared infrastructures. These span the widest range of designs and ages in the review, from fetal MRI (Cook 2024) to a post-trauma adult sample (Webb 2021), and tend to recruit from urban contexts, yielding the more demographically diverse samples in the corpus. Their smaller sample sizes and site-specific recruitment trade statistical power for demographic and geographic specificity. |

*Note.* Sample sizes across included studies ranged from 68 to 9,475 participants (median = 1,824), with the largest samples concentrated among ABCD-derived studies.

### 6. Adolescent Concentration and Lifespan Gaps

The included studies skewed markedly toward adolescent samples, many originating using data obtained from the Adolescent Brain and Cognitive Development (ABCD) Study ^110^, with a secondary cluster of studies examining neonatal and infant populations. This developmental concentration leaves critical gaps across the lifespan, including early and middle childhood, adulthood, and older adulthood, where the neural consequences of cumulative area-level exposure remain largely uncharacterized. The adolescent concentration is in part an outcome of the field’s use of the ABCD Study, a large, multi-site longitudinal dataset that has become the dominant infrastructure for area-level neuroscience research in the United States. While the ABCD Study offers unparalleled statistical power and geographic coverage, its demographic profile and recruitment strategy shape the questions that can be asked and the populations whose experiences are represented. Over-reliance on a single dataset risks conflating dataset-specific findings with general principles, and limits the identification of effects that may be specific to other developmental periods or population contexts. One notable opportunity to address the lower end of this gap is the HEALthy Brain and Child Development (HBCD) Study, a multi-site prospective cohort examining brain and behavioral development from the prenatal period through age 10 ^111,112^. A central aim of HBCD is to investigate associations between adverse environments and socioeconomic disadvantage and the development of brain and behavior across early childhood. The mechanisms through which area-level neighborhood context may shape functional brain organization are likely to differ across development, though direct empirical comparisons across developmental periods remain limited. Early childhood represents a sensitive period for the development of corticolimbic circuitry supporting multiple processes, including but not limited to threat and stress regulation ^31,113,114^. Adolescence involves pubertal reorganization of frontolimbic and default mode networks ^115^; adulthood is characterized by the accumulation of chronic stress exposures; and older adulthood may see differential effects of lifetime area-level disadvantage on age-related trajectories of brain function. A lifespan perspective on the neuroscience of area-level factors requires investigation in understudied age groups, particularly early childhood and older adulthood, using designs capable of examining cumulative exposure across multiple developmental stages.

### 7. Opportunities for Large-Scale Network Analysis and Precision Functional Mapping

Beyond addressing these gaps, several methodological frontiers offer promise for advancing neighborhood neuroscience. The first is the broader adoption of large-scale functional network analysis, including graph-theoretic approaches described above, as well as data-driven methods such as independent component analysis (ICA) and community detection algorithms that do not impose a priori network boundaries. These approaches may be better suited to detecting how area-level context shapes the global architecture of functional brain organization, rather than its expression within any single network or connection. The second opportunity is the application of precision functional mapping (PFM) approaches to questions of area-level neuroscience. Precision functional mapping, exemplified by dense-sampling paradigms such as the Midnight Scan Club ^116^, characterizes functional brain organization at the level of the individual rather than the group. While dense-sampling paradigms require more data per participant, methods designed to operate on more limited data, including template matching ^117^, multi-session hierarchical Bayesian modeling ^118^, and non-negative matrix factorization, make person-specific network estimation tractable in existing datasets such as ABCD. Critically, standard atlas-based and group-averaged parcellation schemes impose network boundaries derived primarily from samples of convenience, typically healthy, white, university-affiliated adults, that may not accurately reflect the functional network organization of individuals from different environmental backgrounds ^119^ or age groups. If neighborhood disadvantage systematically shifts the boundaries, strength, or topology of intrinsic functional networks, then group-averaged parcellations may obscure precisely the individual differences that are of scientific and clinical interest. Applying individualized parcellation approaches to samples with rich neighborhood-level data would enable researchers to ask whether and how place-based neighborhood context impacts the very architecture of a person’s functional connectome.

## Implications and Future Directions

Together, the associations between area-level disadvantage and functional brain connectivity have been observed across a growing number of studies; enough to warrant continued investigations into how area-level context is linked to functional brain organization, and heterogeneous enough in measurement, analytic approach, and sample composition to resist strong conclusions about mechanisms, developmental specificity, or generalizability. Characterizing how area-level factors may become biologically embedded in functional brain organization can, over time, sharpen the mechanistic and developmental understanding that such efforts draw on, especially clarifying when area-level exposures exert their greatest influence and through which neural systems. We are cautious not to overstate what current evidence supports, however. The case for place-based investment, e.g., housing, schools, violence reduction, environmental remediation, rests on well-established social and economic grounds. We provide a non-exhaustive list to attend to opportunities for progress; convergence on validated, geocoded area-level measures, adoption of network-level and precision analytic approaches, transparent reporting of sample demographics at every stage of the research pipeline, and investment in understudied developmental periods and populations. The growing field of understanding area-level factors in neuroscience depends on ensuring that the methods used to study the brain are as rigorous, representative, and conscientious as the questions that motivate the work.

## Data availability

The completed data-charting table containing all extracted variables (sample characteristics, area-level measure and its operationalization, geographic resolution, functional-connectivity approach, and coded direction of association) for the 28 included studies is publicly available in the study’s Open Science Framework repository (https://doi.org/10.17605/OSF.IO/S7VE6).

## CRediT authorship contribution statement

**JAR:** Conceptualization, Methodology, Investigation, Data curation, Formal analysis, Visualization, Project administration, Writing – original draft, Writing – review & editing. **CA:** Formal analysis, Writing – review & editing. **GR:** Formal analysis, Writing – review & editing. **EG:** Formal analysis, Writing – review & editing. **JR:** Formal analysis, Writing – review & editing. **GG:** Formal analysis, Writing – review & editing. **MEL:** Supervision, Writing – review & editing. **RP:** Conceptualization, Supervision, Writing – review & editing.

## Acknowledgments

(JAR): Stanford University Knight-Hennessy Scholars Program, National Academies of Sciences, Engineering, and Medicine’s Ford Foundation Predoctoral Fellowship, Institute of International Education Quad Fellowship, and the National Science Foundation’s Graduate Research Fellowship Program. (EG): National Science Foundation’s Graduate Research Fellowship Program. (GR): Stanford University Knight-Hennessy Scholars Program, Institute of International Education Quad Fellowship, and a Bill and Melinda Gates Millennium Scholarship. (JR): Postdoctoral Award for Collaborative Excellence, Yale Kavli Institute for Neuroscience. The authors would also like to thank Amanda Woodward and Stanford University School of Medicine’s Lane Medical Library for their assistance with the scoping review.

