## Supplemental Materials for "Area-level context and functional brain organization across the lifespan"

**Author(s)**

Jocelyn A. Ricard^1^, Chase Antonacci^1^, Gabriel Reyes^1^, Eugenia Giampetruzzi^1^, Jivesh Ramduny^2^, Gracie Grimsrud^1^, Monica Ellwood-Lowe^1^, Russell A. Poldrack^1^

1. Stanford University, Stanford, CA United States
2. Yale University, New Haven, CT United States

**Corresponding Author:**

Jocelyn A. Ricard,

450 Jane Stanford Way, Building 420

Stanford University

Stanford, CA 94305

**Table S1. Characteristics of self-report studies**

| **Source** | **1st-Author Affiliation** | **N** | **Unit of Analysis** | **Area-Level Measure** |
| --- | --- | --- | --- | --- |
| Butler et al, 2024^1^ | United States | 233 | Self-report | Exposure to violence |
| Goetschius et al, 2020^2^ | United States | 175 | Self-report | Exposure to violence |
| Keding et al, 2025^3^ | United States | 1133 | Self-report | Exposure to violence |
| Mattheiss et al, 2022^4^ | United States | 46 | Self-report | Exposure to violence |
| Reda et al, 2021^5^ | United States | 52 | Self-report | Exposure to violence |
| Saxbe et al, 2018^6^ | United States | 22 | Self-report | Exposure to violence |

**Supplementary Table S1. Studies using self- or informant-reported neighborhood measures (excluded from the main synthesis; k = 6).** Characteristics of the six resting-state fMRI studies that assessed functional connectivity in relation to self- or informant-reported neighborhood constructs (e.g., exposure to community violence) rather than geocoded, validated area-level indices, and were therefore excluded from the main synthesis. Columns report first-author and year, country of first-author affiliation, sample size, unit of analysis, and the self-report measure used. Cataloged to document the broader landscape and support future work comparing self-reported and geocoded operationalizations of area-level context.

**Supplementary Methods.** Full electronic search strategy. Complete search strings as implemented in all four databases (PubMed, Embase, Scopus, and Web of Science), including controlled-vocabulary (MeSH/Emtree) and free-text terms across the three conceptual domains searched; (1) brain and functional connectivity, (2) area-level/neighborhood characteristics, and (3) validated composite area-level deprivation indices with Boolean structure, per-database set counts, publication-type exclusions applied at the search stage, and search dates (6–7 October 2025).

**Full search:**

(("Brain"[MeSH Terms] OR "Brain"[Text Word] OR "neural"[Text Word] OR "neuro*"[Text Word] OR "functional connectivity"[Text Word] OR "functional networks"[Text Word] OR "functional network"[Text Word] OR "resting state"[Text Word] OR "network connectivity"[Text Word]) AND ("Neighborhood Characteristics"[MeSH Terms] OR "Exposure to Violence"[MeSH Terms] OR "Neighborhood Characteristics"[Text Word] OR "neighborhood threat"[Text Word] OR "Neighborhood deprivation"[Text Word] OR "neighborhood disadvantage"[Title/Abstract:~3] OR "neighborhood disadvantages"[Title/Abstract:~3] OR "neighborhood disadvantaged"[Title/Abstract:~3] OR "neighborhood safety"[Text Word] OR "neighborhood effect"[Text Word] OR "neighborhood effects"[Text Word] OR "neighborhood environment"[Text Word] OR "area deprivation index"[Text Word] OR "child opportunity index"[Text Word] OR "socioeconomic status index"[Text Word] OR "social vulnerability index"[Text Word] OR "distressed communities index"[Text Word] OR "social deprivation index"[Text Word] OR "census tract-level socioeconomic status"[Text Word] OR "Townsend deprivation index"[Text Word] OR "material community deprivation index"[Text Word] OR "neighborhood socioeconomic status"[Text Word] OR "community need index"[Text Word] OR "national neighborhood data archive"[Text Word] OR "exposure to crime"[Text Word] OR "crime rate"[Text Word] OR "opportunity atlas"[Text Word] OR "community violence"[Text Word]) AND "English"[Language]) NOT ("Editorial"[Publication Type] OR "Review"[Publication Type])

**Pubmed search:**

Final PubMed Search:

Date searched: 10/6/25

| **Set #** | **Concepts** | **Syntax** | **Results** |
| --- | --- | --- | --- |
| 1 | Brain and neural function | "Brain"[MeSH Terms] OR "Brain"[Text Word] OR "neural"[Text Word] OR "neuro*"[Text Word] OR "functional connectivity"[Text Word] OR "functional networks"[Text Word] OR "functional network"[Text Word] OR "resting state"[Text Word] OR "network connectivity"[Text Word] | 4,206,266 |
| 2 | Neighborhood disadvantage | "Neighborhood Characteristics"[MeSH Terms] OR "Exposure to Violence"[MeSH Terms] OR "Neighborhood Characteristics"[Text Word] OR "neighborhood threat"[Text Word] OR "Neighborhood deprivation"[Text Word] OR "neighborhood disadvantage"[Title/Abstract:~3] OR "neighborhood disadvantages"[Title/Abstract:~3] OR "neighborhood disadvantaged"[Title/Abstract:~3] OR "neighborhood safety"[Text Word] OR "neighborhood effect"[Text Word] OR "neighborhood effects"[Text Word] OR "neighborhood environment"[Text Word] OR "area deprivation index"[Text Word] OR "child opportunity index"[Text Word] OR "socioeconomic status index"[Text Word] OR "social vulnerability index"[Text Word] OR "distressed communities index"[Text Word] OR "social deprivation index"[Text Word] OR "census tract-level socioeconomic status"[Text Word] OR "Townsend deprivation index"[Text Word] OR "material community deprivation index"[Text Word] OR "neighborhood socioeconomic status"[Text Word] OR "community need index"[Text Word] OR "national neighborhood data archive"[Text Word] OR "exposure to crime"[Text Word] OR "crime rate"[Text Word] OR "opportunity atlas"[Text Word] OR "community violence"[Text Word] | 12,966 |
| 3 | combining | #1 AND #2 | 730 |
| 4 | Limits: English language, remove editorials and reviews | #3 AND English[language] NOT ("Editorial" [Publication Type] OR "Review" [Publication Type]) | 683 |

Full Search:

(("Brain"[MeSH Terms] OR "Brain"[Text Word] OR "neural"[Text Word] OR "neuro*"[Text Word] OR "functional connectivity"[Text Word] OR "functional networks"[Text Word] OR "functional network"[Text Word] OR "resting state"[Text Word] OR "network connectivity"[Text Word]) AND ("Neighborhood Characteristics"[MeSH Terms] OR "Exposure to Violence"[MeSH Terms] OR "Neighborhood Characteristics"[Text Word] OR "neighborhood threat"[Text Word] OR "Neighborhood deprivation"[Text Word] OR "neighborhood disadvantage"[Title/Abstract:~3] OR "neighborhood disadvantages"[Title/Abstract:~3] OR "neighborhood disadvantaged"[Title/Abstract:~3] OR "neighborhood safety"[Text Word] OR "neighborhood effect"[Text Word] OR "neighborhood effects"[Text Word] OR "neighborhood environment"[Text Word] OR "area deprivation index"[Text Word] OR "child opportunity index"[Text Word] OR "socioeconomic status index"[Text Word] OR "social vulnerability index"[Text Word] OR "distressed communities index"[Text Word] OR "social deprivation index"[Text Word] OR "census tract-level socioeconomic status"[Text Word] OR "Townsend deprivation index"[Text Word] OR "material community deprivation index"[Text Word] OR "neighborhood socioeconomic status"[Text Word] OR "community need index"[Text Word] OR "national neighborhood data archive"[Text Word] OR "exposure to crime"[Text Word] OR "crime rate"[Text Word] OR "opportunity atlas"[Text Word] OR "community violence"[Text Word]) AND "English"[Language]) NOT ("Editorial"[Publication Type] OR "Review"[Publication Type])

**Final Embase Search:**

Date searched: 10/6/25

| **Set #** | **Concepts** | **Syntax** | **Results** |
| --- | --- | --- | --- |
| 1 | Brain and neural function | 'brain'/exp OR brain:ti,ab,kw OR neural:ti,ab,kw OR neuro*:ti,ab,kw OR 'functional connectivity':ti,ab,kw OR 'functional networks':ti,ab,kw OR 'functional network':ti,ab,kw OR 'resting state':ti,ab,kw OR 'network connectivity':ti,ab,kw | 5,884,375 |
| 2 | Neighborhood disadvantage | 'neighborhood characteristic'/exp OR 'exposure to violence'/exp OR 'Neighborhood Characteristics':ti,ab,kw OR 'neighborhood threat':ti,ab,kw OR  'Neighborhood deprivation':ti,ab,kw OR ('neighborhood' NEAR/3 'disadvantage') OR ('neighborhood' NEAR/3 'disadvantages') OR ('neighborhood' NEAR/3 'disadvantaged') OR 'neighborhood safety':ti,ab,kw OR 'neighborhood effect':ti,ab,kw OR 'neighborhood effects':ti,ab,kw OR 'neighborhood environment':ti,ab,kw OR 'area deprivation index':ti,ab,kw  OR 'child opportunity index':ti,ab,kw  OR 'socioeconomic status index':ti,ab,kw OR 'social vulnerability index':ti,ab,kw  OR 'distressed communities index':ti,ab,kw OR 'social deprivation index':ti,ab,kw  OR 'census tract-level socioeconomic status':ti,ab,kw  OR 'Townsend deprivation index':ti,ab,kw  OR 'material community deprivation index':ti,ab,kw  OR 'neighborhood socioeconomic status':ti,ab,kw  OR 'community need index':ti,ab,kw  OR 'national neighborhood data archive':ti,ab,kw  OR 'exposure to crime':ti,ab,kw OR 'crime rate':ti,ab,kw OR 'opportunity atlas':ti,ab,kw  OR 'community violence':ti,ab,kw | 18,207 |
| 3 | combining | #1 AND #2 | 1,451 |
| 4 | Limits: English language, remove editorials and reviews | #3 AND [english]/lim NOT ('conference abstract'/it OR 'conference review'/it OR 'editorial'/it OR 'review'/it) | 911 |

**Final Scopus Search:**

Date searched: 10/6/25

| **Set #** | **Concepts** | **Syntax** | **Results** |
| --- | --- | --- | --- |
| 1 | Brain and neural function | TITLE-ABS-KEY(brain OR neural OR neuro* OR "functional connectivity" OR "functional networks" OR “functional network” OR "resting state" OR "network connectivity") | 6,753,918 |
| 2 | Neighborhood disadvantage | TITLE-ABS-KEY("Neighborhood Characteristics" OR “neighborhood threat” OR “Neighborhood deprivation” OR “neighborhood safety” OR “neighborhood effect” OR “neighborhood effects” OR "neighborhood environment” OR “area deprivation index” OR “child opportunity index” OR “socioeconomic status index” OR “social vulnerability index” OR “distressed communities index” OR “social deprivation index” OR “census tract-level socioeconomic status” OR “Townsend deprivation index” OR “material community deprivation index” OR “neighborhood socioeconomic status” OR “community need index” OR “national neighborhood data archive” OR “exposure to crime” OR “crime rate” OR “'opportunity atlas” OR “community violence”) OR (neighborhood NEAR/3 disadvantage) OR (neighborhood NEAR/3 disadvantages) OR (neighborhood NEAR/3 disadvantaged) | 29,882 |
| 3 | combining | #1 AND #2 | 1,416 |
| 4 | Limits: English language, remove editorials and reviews | #3 limited to English and article | 1,082 |

**Final Web of Science Search:**

Date searched: 10/7/25

| **Set #** | **Concepts** | **Syntax** | **Results** |
| --- | --- | --- | --- |
| 1 | Brain and neural function | TS=(brain OR neural OR neuro* OR "functional connectivity" OR "functional networks" OR “functional network” OR "resting state" OR "network connectivity") | 4,189,769 |
| 2 | Neighborhood disadvantage | TS=("Neighborhood Characteristics" OR “neighborhood threat” OR “Neighborhood deprivation” OR “neighborhood safety” OR “neighborhood effect” OR “neighborhood effects” OR "neighborhood environment” OR “area deprivation index” OR “child opportunity index” OR “socioeconomic status index” OR “social vulnerability index” OR “distressed communities index” OR “social deprivation index” OR “census tract-level socioeconomic status” OR “Townsend deprivation index” OR “material community deprivation index” OR “neighborhood socioeconomic status” OR “community need index” OR “national neighborhood data archive” OR “exposure to crime” OR “crime rate” OR “'opportunity atlas” OR “community violence”) OR (neighborhood NEAR/3 disadvantage) OR (neighborhood NEAR/3 disadvantages) OR (neighborhood NEAR/3 disadvantaged) | 11,763 |
| 3 | combining | #1 AND #2 | 633 |
| 4 | Limits: English language, remove editorials and reviews | #3 limited to English and article | 565 |
